# The PIN1-p38-CtIP signaling axis protects stalled replication forks from deleterious degradation

**DOI:** 10.1101/2024.06.25.600562

**Authors:** Francesca Vivalda, Marco Gatti, Letizia Manfredi, Hülya Dogan, Antonio Porro, Giulio Collotta, Ilaria Ceppi, Christine von Aesch, Vanessa van Ackeren, Sebastian Wild, Martin Steger, Begoña Canovas, Monica Cubillos-Rojas, Antoni Riera, Petr Cejka, Angel R. Nebreda, Diego Dibitetto, Sven Rottenberg, Alessandro A. Sartori

**Affiliations:** Institute of Molecular Cancer Research, University of Zurich, Zurich, Switzerland; Institute of Animal Pathology and Bern Center for Precision Medicine, University of Bern, Bern, Switzerland; Institute for Research in Biomedicine, Università della Svizzera italiana, Bellinzona, Switzerland; NEOsphere Biotechnologies, Martinsried, Germany; Institute for Research in Biomedicine (IRB Barcelona), The Barcelona Institute of Science and Technology, Barcelona, Spain; Departament de Química Inorgànica i Orgànica, Secció de Química Orgànica, Universitat de Barcelona, Barcelona, Spain; ICREA, Barcelona, Spain; Department of Experimental Oncology, Istituto di Ricerche Farmacologiche Mario Negri IRCCS, Milan, Italy; Cancer Therapy Resistance Cluster, Department for Biomedical Research, University of Bern, Bern, Switzerland

## Abstract

Human CtIP plays a critical role in homologous recombination (HR) by promoting the resection of DNA double-strand breaks. Moreover, CtIP maintains genome stability through protecting stalled replication forks from nucleolytic degradation. However, the upstream signaling mechanisms governing the molecular switch between these two CtIP-dependent processes remain largely elusive. Here, we show that phosphorylation of CtIP by the p38α stress kinase and subsequent PIN1-mediated CtIP *cis*-to-*trans* isomerization is required for fork stabilization but dispensable for HR. We found that stalled forks are degraded in cells expressing non-phosphorylatable CtIP or lacking PIN1-p38α activity, while expression of a CtIP *trans*-locked mutant overcomes the requirement for PIN1-p38α in fork protection. We further reveal that *Brca1*-deficient mammary tumor cells that have acquired PARPi resistance regain chemosensitivity after PIN1 or p38α inhibition. Collectively, our findings identify the PIN1-p38-CtIP signaling pathway as a critical regulator of replication fork integrity.

## INTRODUCTION

The maintenance of genome stability relies on the accurate completion of DNA replication during S-phase. The progression of replication forks can be impeded by many internal and external events such as DNA damage and depletion of nucleotide precursors, causing the accumulation of single-stranded DNA and triggering replication stress, a crucial vulnerability of cancer cells^1^. Numerous factors have been implicated in the protection and recovery of stalled replication forks to prevent their collapse into highly mutagenic DNA double-strand breaks (DSBs)^2^. This includes proteins involved in homologous recombination (HR), most notably BRCA1 and BRCA2, which protect nascent DNA from degradation by the MRE11 exonuclease^3^. Moreover, we have recently uncovered a key role for CtIP in the protection of stalled forks from nucleolytic degradation by DNA2, through a mechanism that acts independently from its well-established DSB end resection function^4,5^. Accordingly, CtIP-T847A and -S327A phosphomutants defective in MRE11-RAD50-NBS1 (MRN) and BRCA1 interaction^6–8^, respectively, and thus, in HR, are proficient in fork protection. In contrast, CtIP nuclease-defective mutants proficient in DNA-end resection and HR were shown to cause fork degradation upon replicative stress^4^. However, the regulatory mechanisms mediating CtIP’s role in fork protection have remained largely elusive.

PIN1 is a unique phosphorylation-specific peptidyl-prolyl *cis*-to-*trans* isomerase reported to act as a molecular switch and pivotal modulator of multiple cellular processes. Consistently, aberrant PIN1 activity has been linked to a plethora of human pathologies, including cancer and neurodegeneration^9,10^. Through proteomics, we previously identified several prominent DNA damage response factors as potential PIN1 interaction partners, including BRCA1 and CtIP^11^. We further demonstrated that PIN1 can bind to two conserved phosphorylated S/T-P motifs (pS276 and pT315) but preferentially isomerizes the pS276-P277 peptide bond in CtIP. This conformational change ultimately regulates CtIP protein turn-over, thereby fine-tuning DNA-end resection^11^. While we could reveal CDK2 as the major kinase responsible for CtIP-T315 phosphorylation during S and G2 phase, the proline-dependent kinase phosphorylating CtIP at S276 has not yet been identified. In addition to the canonical ATM and ATR signaling pathways, the stress-induced p38 mitogen-activated kinase (p38 MAPK) family has been reported to contribute to cell cycle arrest in response to genotoxic agents^12,13^. Notably, p38α, the best characterized and ubiquitously expressed isoform of the p38 MAPK family, was reported to restrain chromosome instability in mammary tumor cells and to phosphorylate several S/T-P motifs in recombinant CtIP^14^.

Here, we report that CtIP phosphorylation by p38α kinase at S276 followed by PIN1-mediated *cis*-to-*trans* isomerization of the pS276/P277 peptide bond is necessary for the protection of stalled forks from nucleolytic degradation, but dispensable for HR. Expression of CtIP-S276A or inhibition of PIN1/p38α activity trigger forks degradation. We find that PIN1/p38α-mediated CtIP isomerization is critical for CtIP accumulation at stalled forks. Finally, we reveal that *Brca1*-deficient mammary tumor cells, that acquired resistance to PARP inhibitor via CtIP-dependent restoration of fork stability, regain chemosensitivity after PIN1 or p38α inhibition. Collectively, we define the p38α-PIN1-CtIP phosphorylation-isomerization cascade as a crucial regulatory mechanism preserving replication fork integrity.

## RESULTS

### Isomerization of the phospho-S276-P277 motif in CtIP protects stalled forks from nucleolytic degradation

In our past studies, we have identified CtIP as a target of PIN1 isomerization and demonstrated that CtIP depletion triggers DNA2-dependent fork degradation in a BRCA1-independent manner^4,11^. However, the mechanism underlying CtIP-mediated fork protection remained unknown.

This prompted us to investigate the potential role of CtIP isomerization in fork protection. We first performed DNA fiber spreading assays upon treatment with the ribonucleotide reductase inhibitor and replication stalling agent hydroxyurea (HU) to measure fork degradation in cells stably expressing different siRNA-resistant GFP-CtIP variants, established previously^11^. We observed that, unlike wild-type (wt) CtIP, phosphomutants defective in PIN1 binding (T315A), isomerization (S276A) or both (S276A/T315A) failed to restore fork stability (Figure 1A). Moreover, temporary inhibition of DNA2 nuclease activity as well as depletion of the SMARCAL1 DNA translocase alleviated degradation of HU-stalled forks in cells expressing isomerization-defective CtIP mutants (Figures 1B, 1C and S1A). These results are consistent with our previous findings showing that CtIP prevents extensive nascent strand degradation by DNA2 after fork reversal^4^ and suggest that PIN1-CtIP interaction is required to maintain fork stability in cells experiencing replication stress. We next wanted to assess more directly whether *cis* to *trans* prolyl-peptide bond isomerization at the pS276-P277 motif in CtIP is critical for replication fork protection. Therefore, we generated U2OS cells inducibly expressing GFP-tagged *trans*-locked mutants of CtIP, in which P277 was substituted with alanine, either alone (P277A) or in combination with S276A (S276A/P277A). First, we analyzed CtIP-S276 phosphorylation and CtIP-PIN1 interaction in cells expressing P277A mutants and found that both events are strongly impaired (Figures S1B and S1C), indicating a critical role for P277 in S276 phosphorylation, and, consequently, in PIN1 binding. Remarkably, however, employing two alternative experimental approaches to analyze fork stability, expression of CtIP *trans*-locked variants (P277A or S276A/P277A) rescued fork degradation in CtIP-depleted cells, indicating that forced *trans*-geometry of the P277 peptide bond can compensate for the lack of S276 phosphorylation and PIN1 binding (Figure 1D and S1D). Notably, the observed differences in fork stability between CtIP mutant cells could not be attributed to differences in CtIP protein stability (Figure S1E).

**Figure 1.**
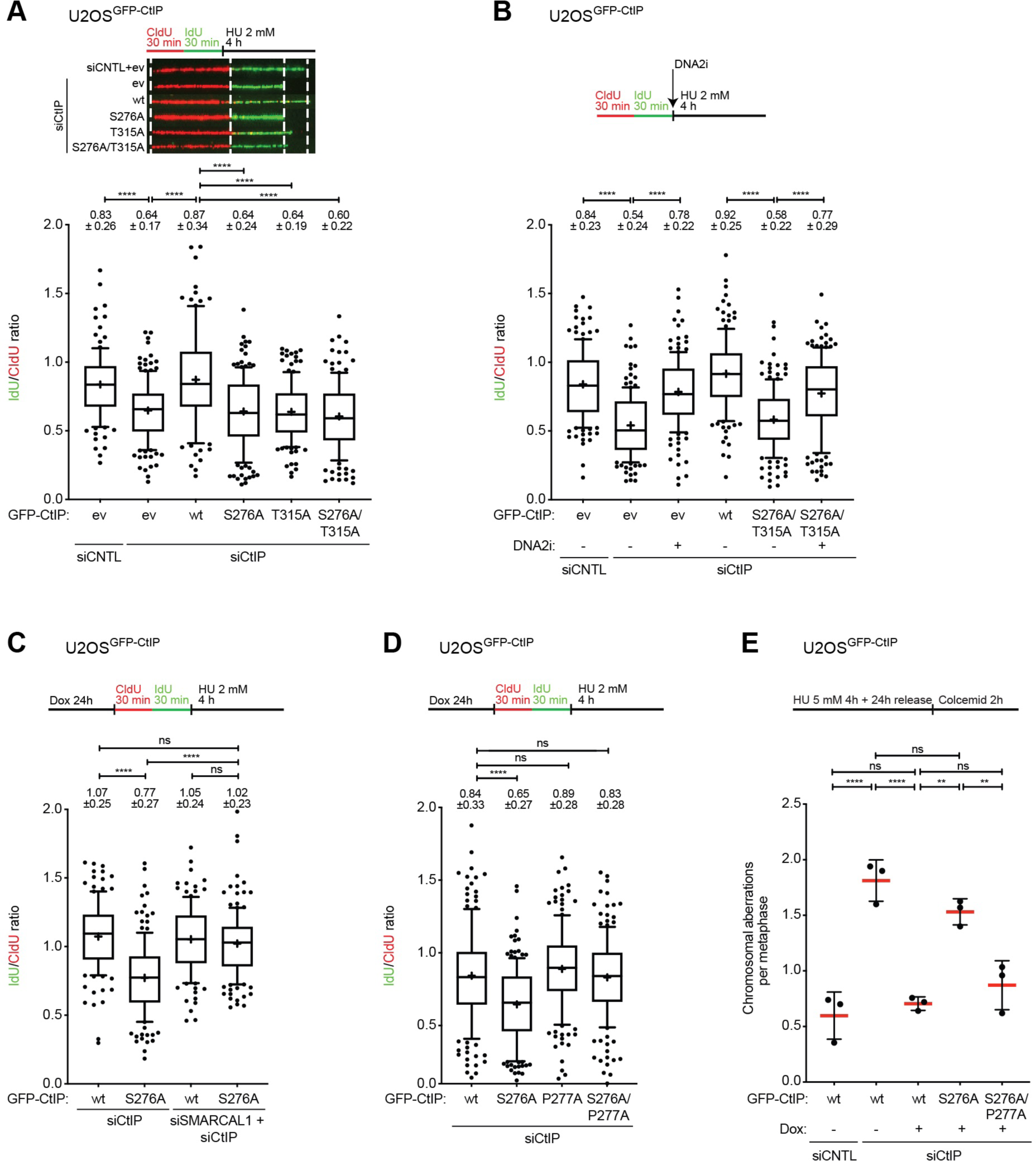
CtIP *cis*-to-*trans* isomerization protects stalled forks from nucleolytic degradation. **(A)** Fork degradation was evaluated upon HU treatment in U2OS cells depleted of endogenous CtIP and stably expressing either GFP empty vector (ev), or siCtIP-resistant GFP-CtIP wild-type (wt), S276A, T315A, and S276A/T315A variants. Representative DNA fiber images are shown (top). **(B)** Fork degradation was evaluated upon HU treatment in U2OS cells depleted of endogenous CtIP and stably expressing siCtIP-resistant GFP-CtIP wt or S276A/T315A variants. In addition, cells were either mock-treated or treated with the DNA2 inhibitor NSC-105808 (2 µM, simultaneously with HU). **(C)** Fork degradation was evaluated upon HU treatment in U2OS cells inducibly expressing siCtIP-resistant GFP-CtIP wt or S276A variants and depleted of endogenous CtIP alone, or co-depleted of CtIP and SMARCAL1. **(D)** Fork degradation was evaluated upon HU treatment in U2OS cells depleted of endogenous CtIP and inducibly expressing siCtIP-resistant GFP-CtIP wt, S276A, P277A, or S276A/P277A (*trans*-locked) variants. **(A-D)** Box and whisker plots of IdU/CldU-tract length ratios for individual replication forks are shown. Numbers indicated above the individual plots represent the mean ratios ± standard deviation. Schematics of the CldU/IdU pulse-labelling protocol are shown (top). **(E)** Metaphase spread analysis upon HU treatment of U2OS cells depleted of endogenous CtIP and inducibly expressing siCtIP-resistant GFP-CtIP wt, S276A, or S276A/P277A (*trans*-locked) variants. Chromatid breaks, fusions and radials were scored. Total chromosomal aberrations per metaphase are shown. The mean (red line) with standard deviation of biological triplicates is shown.

We have previously demonstrated that engineered U2OS*^Cas9/CtIP^* cells, lacking two out of three existing CtIP gene copies, are largely proficient in resecting DSBs and HR but display replication stress-associated phenotypes comparable to those detected in CtIP-depleted cells, including nascent strand degradation and elevated levels of chromatin-bound RPA following HU treatment^4^. Using quantitative image-based cytometry (QIBC), we observed that stable expression of CtIP-wt in U2OS*^Cas9/CtIP^* cells alleviated HU-induced RPA hyperaccumulation on chromatin, whereas the CtIP-S276A phosphomutant did not. Complementing the hypomorphic cells with the CtIP-S276A/P277A *trans*-locked mutant, however, restored chromatin-bound RPA levels comparable to CtIP-wt cells, indicating that CtIP isomerization limits the accumulation of RPA on single-stranded DNA due to replication stress (Figures S1F and S1G).

Excessive nucleolytic resection of reversed forks hinders the faithful completion of DNA replication during S-phase, which can potentially cause chromosomal aberrations. Consistent with a role for CtIP isomerization in maintaining genome stability in response to replication stress, we detected a significantly higher frequency of HU-induced chromosomal aberrations in CtIP-depleted cells, which was rescued by the expression of CtIP-wt or -S276A/P277A *trans*-locked variants, but not by the S276A phosphomutant (Figures 1E, S1H and S1I). Collectively, our findings implicate *cis*-to-*trans* isomerization of the CtIP pS276-P277 peptide bond as a critical step in preventing the nucleolytic degradation of nascent DNA after replication stress.

### PIN1-catalyzed CtIP isomerization is required for fork protection but dispensable for HR

To further corroborate the contribution of CtIP isomerization by PIN1 in fork stabilization, we made use of KPT-6566, a selective and covalent prolyl isomerase PIN1 inhibitor^15^. We found that 1 hour pretreatment of cells with KPT-6566 induced fork degradation in a dose-dependent manner without affecting CtIP protein stability (Figures S2A and S2B). PIN1-mediated BRCA1 isomerization was previously reported to protect forks from MRE11-dependent degradation^16^. In agreement with that study, we found that combined treatment of cells with KPT-6566 and mirin, an inhibitor of MRE11 3ʹ–5ʹ exonuclease activity^17^, restored fork stability (Figure 2A). Strikingly, co-treatment with NSC-105808, a selective and potent DNA2 nuclease inhibitor^18^, also rescued fork degradation (Figure 2A), indicating that PIN1 activity counteracts both MRE11- and DNA2-mediated resection of regressed DNA arms at stalled forks, and suggesting that both BRCA1 and CtIP functions at stalled forks are regulated by phosphorylation-dependent isomerization. To dissect the specific role of CtIP isomerization in this scenario, we assessed HU-induced fork degradation in cells harboring different CtIP variants and pretreated with the PIN1 inhibitor. Interestingly, expression of the CtIP S276A/P277A *trans*-locked mutant resulted in a significant, yet incomplete restoration of fork stability (Figure 2B), consistent with a scenario in which PIN1 inhibition simultaneously compromises BRCA1- and CtIP-mediated fork protection pathways. Therefore, expression of CtIP S276A/P277A was unable to rescue fork stability in BRCA1-depleted cells (Figures 2C and S2C).

**Figure 2.**
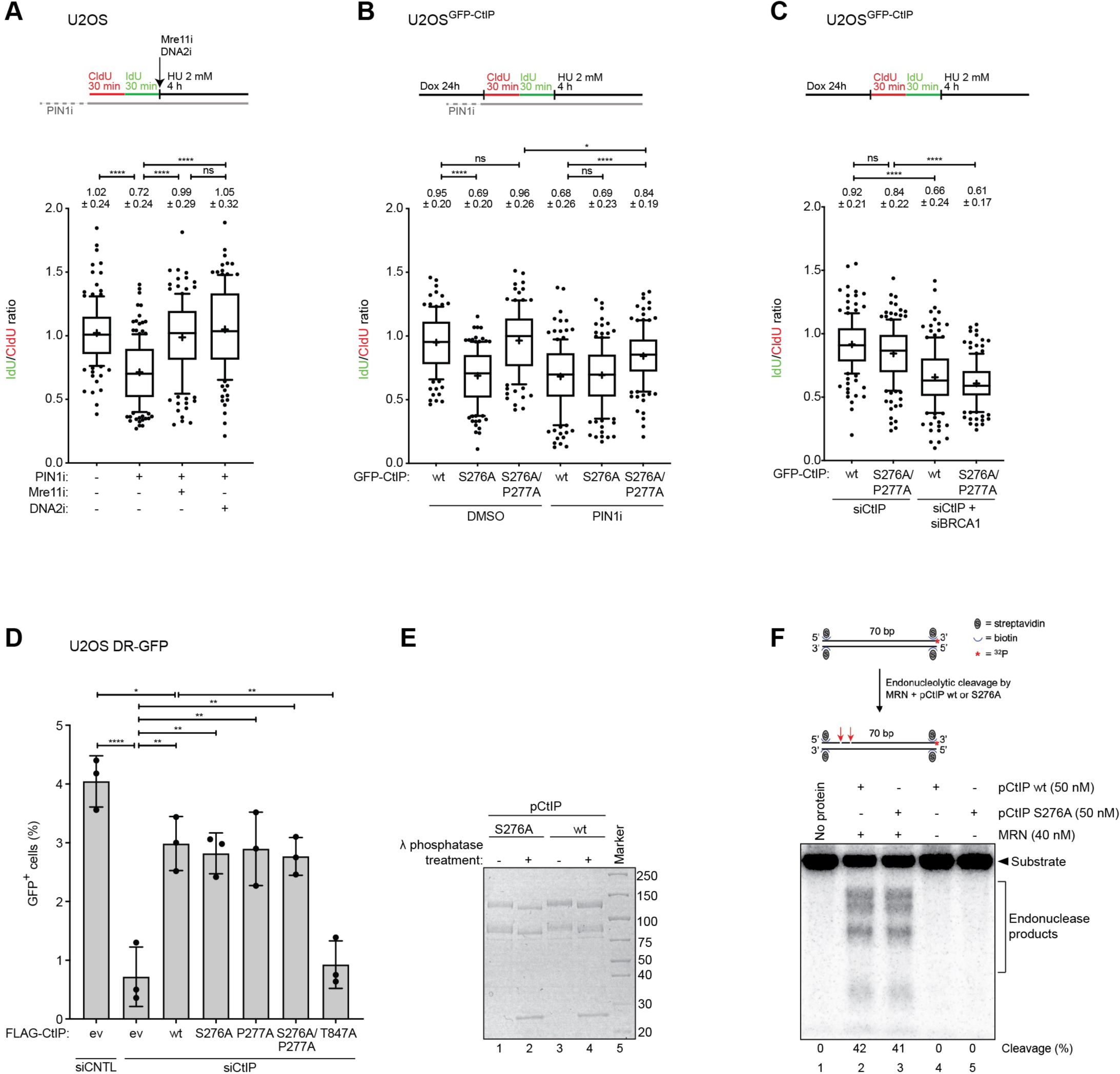
CtIP isomerization by PIN1 promotes fork stability but is dispensable for HR. **(A)** Fork degradation was evaluated upon HU treatment in U2OS cells pre-treated with the PIN1 inhibitor KPT-6566 (10 µM) alone or in combination with either the Mre11 inhibitor Mirin (25 µM) or the DNA2 inhibitor NSC-105808 (2 µM). **(B)** Fork degradation was evaluated upon HU treatment in U2OS cells depleted of endogenous CtIP and inducibly expressing siCtIP-resistant GFP-CtIP wt, S276A or S276A/P277A *trans*-locked mutant. In addition, cells were either mock-treated or treated with the PIN1 inhibitor KPT-6566 (10 µM, 1 h before labelling). **(C)** Fork degradation was evaluated upon HU treatment in U2OS cells inducibly expressing siCtIP-resistant GFP-CtIP wt or S276A/P277A *trans*-locked mutant and depleted of endogenous CtIP alone, or co-depleted of CtIP and BRCA1. **(A-C)** Box and whisker plots of IdU/CldU-tract length ratios for individual replication forks are shown. Numbers indicated above the individual plots represent the mean ratios ± standard deviation. Schematics of the CldU/IdU pulse-labelling protocol are shown (top). **(D)** HR efficiency was evaluated in U2OS DR-GFP cells depleted for endogenous CtIP and transfected with either empty vector (ev) or indicated siCtIP-resistant FLAG-CtIP constructs. Cells were co-transfected with the I-SceI expression plasmid and harvested at 48h post-transfection and analyzed by flow cytometry for GFP signal. Data are shown as percentage of GFP-positive cells. **(E)** Electrophoretic mobility of recombinant CtIP wild-type (wt) and S276A either not-treated or treated with λ phosphatase. **(F)** Endonuclease assay with recombinant MRN complex and either phosphorylated CtIP wt or phosphorylated S276A variant on a 5ʹ end-labelled 70 bp-long double-stranded DNA substrate blocked at both ends with streptavidin. The quantitation (cleavage, %) is an average from three independent experiments. Schematic of the substrate and endonucleolytic cleavage is shown (top).

We have previously shown that PIN1 fine-tunes the balance between HR and non-homologous end-joining (NHEJ) primarily through modulating CtIP turnover via phosphorylation-mediated ubiquitination and subsequent proteasomal degradation^11^. Moreover, Luo *et al*. reported that all-*trans* retinoic acid (ATRA), another PIN1 inhibitor leading to PIN1 degradation, disrupts HR and sensitizes breast cancer cells to PARP inhibition due to decreased BRCA1 protein stability^19^. In agreement with this study, treatment of U2OS/DR-GFP cells with 10 µM of the PIN1 inhibitor KPT-6566 led to a significant defect in HR repair activity (Figure S2D). To examine the direct contribution of CtIP isomerization to HR, we performed DSB repair reporter assays in CtIP-depleted U2OS/DR-GFP cells transfected with siRNA-resistant FLAG-CtIP variants (S2E). Strikingly, we found that HR repair was efficiently and equally rescued by expression of CtIP-wt or any of the CtIP isomerization mutants, but not by expression of a T847A phosphomutant defective in stimulating the MRN endonuclease activity that initiates DNA end resection (Figure 2D)^20^. Moreover, CtIP isomerization mutants, but not the CtIP-S327A phosphomutant^8^, were proficient in BRCA1 interaction (Figure S2F). Finally, expression of CtIP *trans*-locked mutants failed to restore HR deficiency in PIN1-inhibited cells (Figure S2G), suggesting that impaired BRCA1 (but not CtIP) isomerization contributes to HR deficiency caused by PIN1 inhibition.

To further distinguish the specific function of CtIP-S276 phosphorylation-dependent isomerization in promoting fork protection *versus* DNA end resection, we performed *in vitro* nuclease assays with recombinant human CtIP and MRN purified from insect cells^20^. Treatment of CtIP-wt and -S276A with λ-phosphatase resulted in the disappearance of an electrophoretic mobility shift, confirming that both variants exist as phosphorylated forms (pCtIP) after the purification procedure (Figure 2E). Importantly, both pCtIP-wt and pCtIP-S276A stimulated the MRN endonuclease, while lacking any detectable intrinsic nuclease activity (Figure 2F). Together, our findings demonstrate that PIN1-mediated CtIP isomerization at the pS276-P277 motif is critically required for fork protection, but dispensable for DSB resection and HR.

### Stress-activated p38α kinase protects stalled forks from degradation by facilitating CtIP isomerization

We have previously demonstrated that pT315 serves as the major CtIP binding site of PIN1, but that CtIP isomerization by PIN1 exclusively happens at the pS276-P277 motif^11^. In addition, using individual phospho-specific CtIP antibodies, we found that treatment of cells with roscovitine, a non-selective CDK1/2 inhibitor, impaired T315 (but not S276) phosphorylation, suggesting that a different proline-dependent kinase is acting upstream to phosphorylate S276 and facilitate CtIP isomerization in response to replication stress^11^.

We noticed that the region encompassing S276-P277 is highly conserved in mammalian CtIP orthologs and matches the consensus sequence for members of the mitogen-activated protein kinase (MAPK) family, especially that of p38α (encoded by *MAPK14*) (Figure S3A). It has been repeatedly reported that HU treatment induced activation of p38 MAPKs in S-phase synchronized cells, as measured by phosphorylation of p38 itself (at T180/Y182) and of MK2, a bona fide downstream p38 substrate^21–23^. Moreover, p38α was shown to directly phosphorylate CtIP on several S/T-P sites in *in vitro* kinase assays, including S276^14^. These findings prompted us to investigate whether p38α interacts with CtIP and participates in CtIP-S276 phosphorylation following DNA replication stress.

First, we performed Myc-trap pulldowns from HEK293T cells transfected with Myc-p38α and found that p38α associates with CtIP but not with Mre11 (Figure 3A). We confirmed CtIP-p38α interaction using a reciprocal approach, retrieving Myc-p38α via immunoprecipitation of endogenous CtIP from HEK293T lysates (Figure S3B). To investigate the role of p38α in CtIP phosphorylation, we performed an anti-phospho-CtIP (S276) immunoprecipitation in non-synchronized U2OS cells expressing GFP-CtIP wt treated with HU alone or in combination with PH-797804, an ATP-competitive, selective p38α/β kinase inhibitor^24^. Western blotting of the immunoprecipitates revealed a moderate increase in CtIP-S276 phosphorylation upon HU treatment, which was reduced upon concomitant p38α inhibition, both events coinciding with p38α activation levels detected in the input samples (Figure 3B). To corroborate this result, we performed GFP-trap assays in the same cells but this time co-treated with the VHL-based PROTAC compound NR-11c, specifically targeting p38α for proteasomal degradation^25^. Probing the pulldowns with the anti-phospho-CtIP (S276) antibody confirmed that p38α mediates HU-induced phosphorylation of CtIP at S276 (Figure S3C). Notably, both experiments revealed the presence of CtIP-pS276 in absence of HU, suggesting basal S276 phosphorylation by p38α (or another proline-directed kinase) in our engineered U2OS osteosarcoma cell line. In accordance with this, we observed p38α phosphorylation in untreated cells (Figure 3B). This could be explained by GFP-CtIP overexpression upon doxycycline addition already releasing a cellular stress signal that is sufficient for p38α activation even in unperturbed conditions. As CtIP-PIN1 interaction relies in part on S276 phosphorylation (Figure S1C), we reasoned that p38α inhibition should similarly impair PIN1-CtIP interaction. Indeed, GFP-trap assays revealed that the binding of FLAG-CtIP-wt to GFP-PIN1 is reduced in cells treated with the p38α inhibitor, to levels comparable to that of FLAG-CtIP-S276A (Figure 3C).

**Figure 3.**
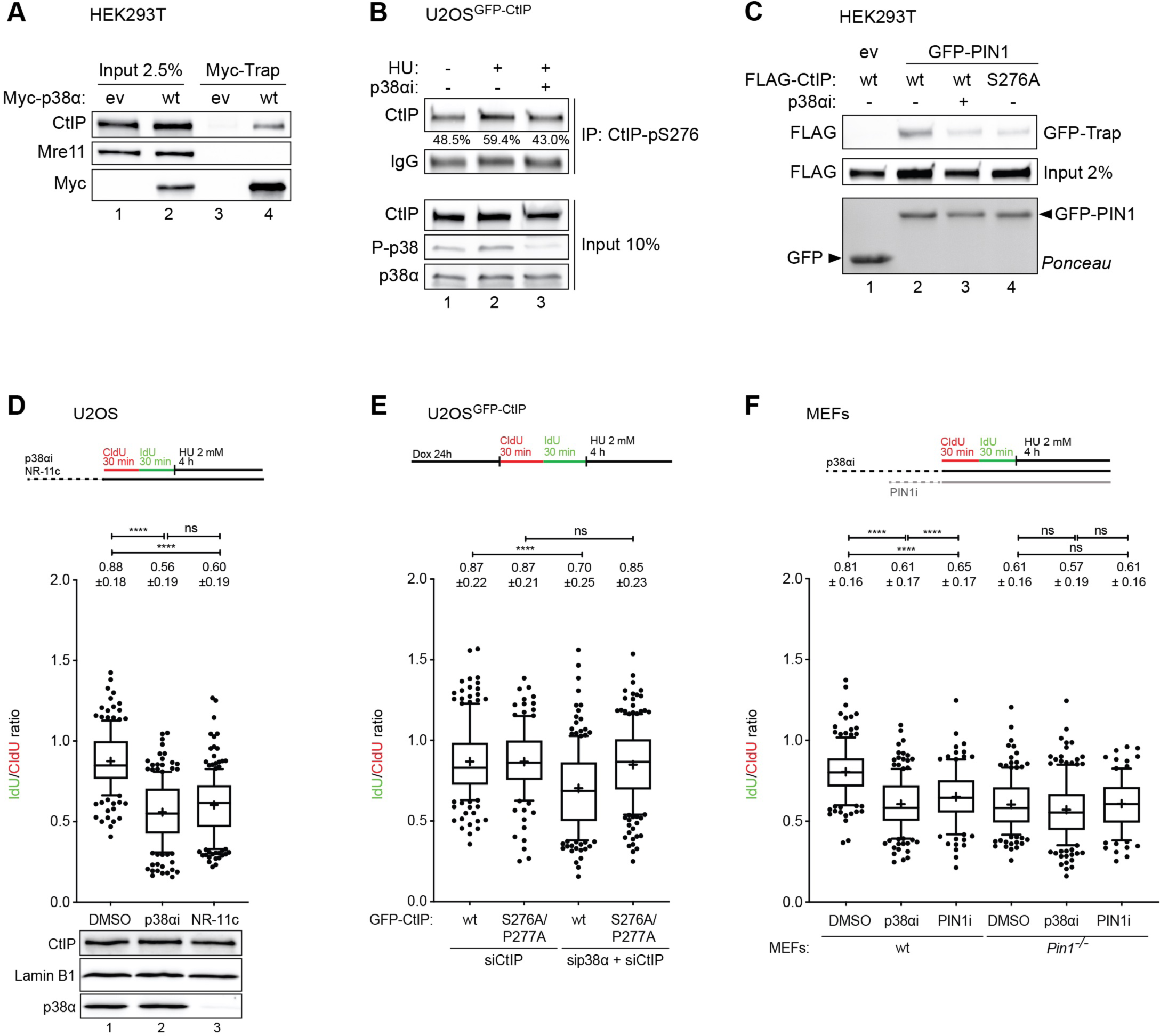
HU-activated p38α phosphorylates CtIP at S276 and facilitates CtIP-dependent fork protection. **(A)** Myc-Trap of HEK293T cells transfected with Myc-p38α. Whole-cell lysates (input) and immunoprecipitates were analyzed by western blotting using specific antibodies. **(B)** Immunoprecipitation (IP) of CtIP-pS276 from U2OS cells inducibly expressing GFP-CtIP either mock-treated or treated with HU (2 mM, 4h). Where indicated, cells were treated with the p38α inhibitor PH-797804 (1 µM, 24h before HU). Whole-cell lysates (input) and immunoprecipitates were analyzed by western blotting using specific antibodies. Densiometric quantification of CtIP band in the IP is shown (% indicates CtIP band intensity vs IgG band intensity). **(C)** GFP-Trap of HEK293T cells co-transfected with GFP-PIN1 and indicated FLAG-CtIP variants. 24h post-transfection, cells were either mock-treated or treated with the p38α inhibitor PH-797804 (1 µM) for 24h. Whole-cell lysates (input) and immunoprecipitates were analyzed by western blotting using specific antibodies. **(D)** Fork degradation was evaluated upon HU treatment in U2OS cells either treated with the p38α inhibitor PH-797804 (1 µM, 24h before HU) or with the p38α PROTAC NR-11c (1 µM, 24h before HU). Western blotting of lysates from the same experiment is shown below. **(E)** Fork degradation was evaluated upon HU treatment in U2OS cells inducibly expressing siCtIP-resistant GFP-CtIP wt or S276A/P277A *trans*-locked mutant and depleted of endogenous CtIP alone, or co-depleted of CtIP and p38α. **(F)** Fork degradation was evaluated upon HU treatment in wild-type mouse embryonic fibroblasts (MEFs) and *Pin1^-/-^* MEFs, pre-treated either for 24h with the p38α inhibitor PH-797804 (1 µM) or for 1h with the PIN1 inhibitor KPT-6566 (10 µM). **(D-F)** Box and whisker plots of IdU/CldU-tract length ratios for individual replication forks are shown. Numbers indicated above the individual plots represent the mean ratios ± standard deviation. Schematics of the CldU/IdU pulse-labelling protocol are shown (top).

It has been previously reported that *Mapk14* deletion (p38αΔ) in mouse mammary tumor cells results in a higher frequency of fork stalling^14^. However, the function of p38α in fork protection during acute HU-induced replication stress has to our knowledge never been elucidated. Thus, we next evaluated fork degradation upon p38α inhibition or degradation. Remarkably, we found that treatment of U2OS cells with either PH-797804 or NR-11c resulted in HU-induced fork degradation (Figure 3D). Moreover, siRNA-mediated depletion of p38α in U2OS resulted in significant fork degradation, consistent with a prominent role of p38α in fork stabilization in response to replication stress (Figure S3D). We earlier showed that expression of the CtIP *trans*-locked mutant resulted in a partial restoration of fork stability in PIN1-inhibited cells (Figure 2B). Remarkably, expressing CtIP-S276A/P277A in p38α-depleted cells fully restored fork protection, indicating a key function of p38α-mediated CtIP phosphorylation in preserving fork integrity upon HU-induced replication stress (Figures 3E and S3E). Consistent with a conserved function of the PIN1-p38α-CtIP signaling axis in fork protection, we found that both PIN1 and p38α inhibition induced fork degradation in spontaneously immortalized primary mouse embryonic fibroblasts (MEFs) (Figures 3F and S3F). Finally, we observed that HU-induced nascent strand degradation in *Pin1* knockout MEFs was not further enhanced upon pre-treatment with the p38α inhibitor, indicating that PIN1 and p38α most likely act in the same fork protection pathway.

### PIN1 and p38α are required for CtIP enrichment at HU-arrested forks

Having established that CtIP *cis*-to-*trans* isomerization at the S276-P277 motif by the collaborative action of p38α and PIN1 is required for fork protection, we next sought to determine whether the conformational change affects CtIP loading and accumulation on stalled replication forks. To test this hypothesis, we employed *in situ* analysis of protein interactions at DNA replication forks (SIRF) using pulsed Edu-CtIP proximity ligation assay (PLA) reactions (CtIP-SIRF)^26,27^. Notably, only PLA signals in cells with comparable EdU intensities were considered to ensure that observed changes in the number of PLA signals/cell did not relate to changes in EdU incorporation between the different experimental conditions. We first performed CtIP-SIRF in unperturbed U2OS cells and could readily detect nuclear PLA signals (Figure 4A), indicating CtIP loading nearby normal ongoing replication forks, as previously reported^28^. After HU treatment, we observed a significant increase in PLA signals, indicating CtIP accumulation at stalled forks (Figure 4A). Interestingly, CtIP loading to stalled forks was impaired when cells were pre-treated with the PIN1 inhibitor prior to addition of HU (Figure 4A). Similar results for CtIP-SIRF in response to HU are obtained when U2OS cells were pre-treated with the p38α inhibitor (Figure 4B), substantiating the importance of PIN1-p38α signaling in facilitating the assembly of CtIP at stalled replication forks. Lastly, we performed SIRF assays in HU-treated cells expressing GFP-tagged CtIP variants (Figure S4A) and observed that the S276A phosphomutant displayed significantly reduced numbers of PLA foci, which were restored to wild-type levels in the S276A/P277A *trans*-locked mutant (Figure 4C). To rule out a more general role of CtIP isomerization in the recruitment to sites of DNA damage, we monitored GFP-CtIP accumulation at microlaser-induced DSBs. However, we could not observe any major differences in the assembly of GFP-CtIP at DSBs in cells either treated with the PIN1 inhibitor or expressing the isomerization-defective CtIP mutants (Figures S4B and S4C). Together, our data reveal a prominent role for PIN1-p38α-mediated CtIP isomerization in efficient loading of CtIP at sites of stalled DNA replication.

**Figure 4.**
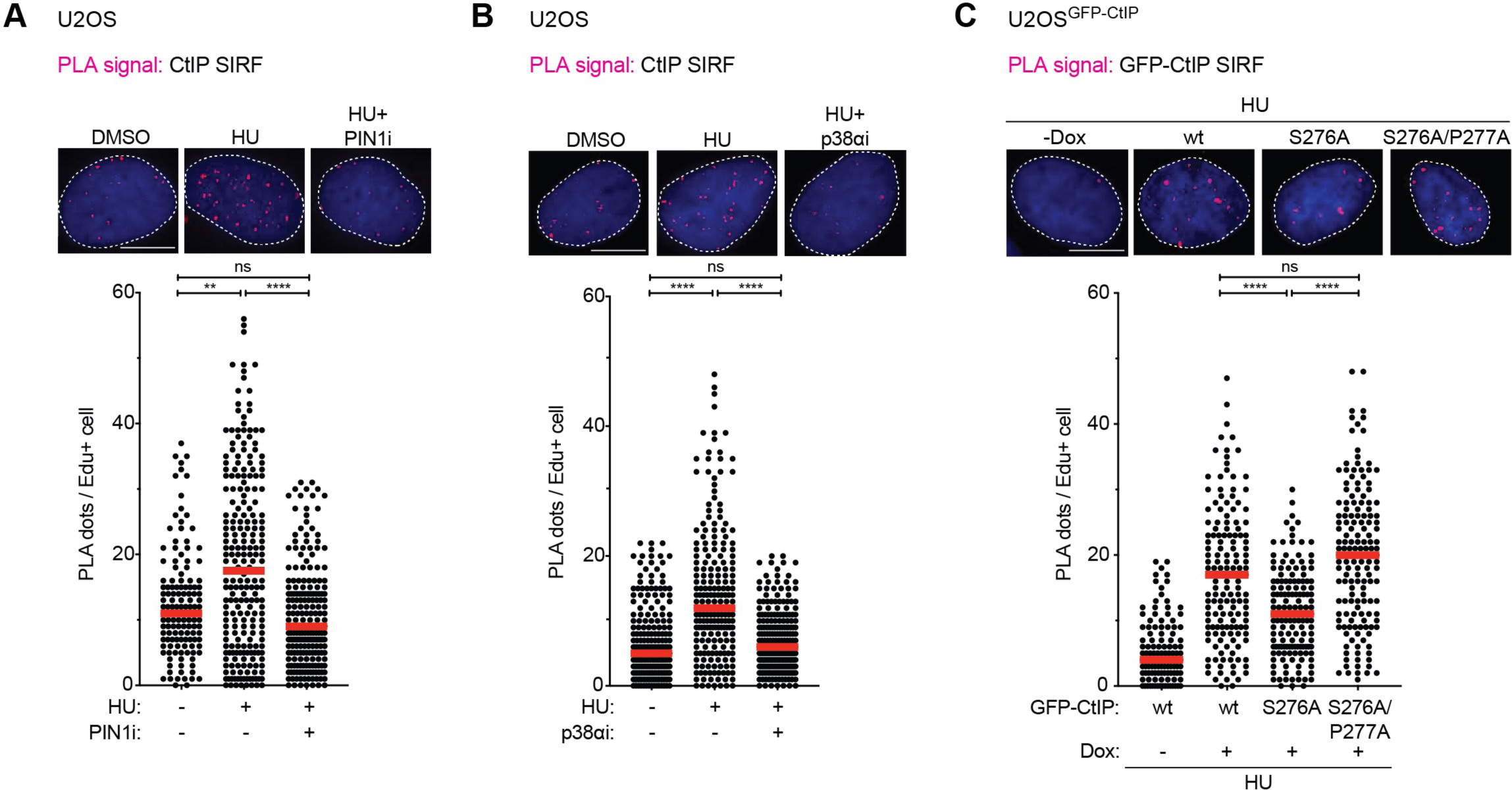
PIN1 and p38α activities are required for CtIP accumulation at stalled replication forks. **(A)** CtIP SIRF assay in U2OS cells pulsed-labelled with EdU (25 µM) for 10 min followed by treatment with HU (2mM) for 4h. Where indicated cells were treated with the PIN1 inhibitor KPT-6566 (10 µM, 1h before EdU labelling). **(B)** CtIP SIRF assay in U2OS cells pulsed-labelled with EdU (25 µM) for 10 min followed by treatment with HU (2mM) for 4h. Where indicated cells were treated with the p38α inhibitor PH-797804 (1 µM, 24h before EdU labelling). **(C)** GFP-CtIP SIRF assay in U2OS cells inducibly expressing siCtIP-resistant GFP-CtIP wt, S276A or S276A/P277A *trans*-locked mutant and depleted of endogenous CtIP. Cells were pulsed-labelled with EdU (25 µM) for 10 min followed by treatment with HU (2mM) for 4h. **(A-C)** Dot plots show the number of PLA foci and the median from at least 120 EdU-positive cells. Representative images are shown on top of each figure. Scale bars, 10 μm.

### Inhibition of PIN1 or p38α overcomes chemoresistance in *Brca1*-deficient mammary tumor cells

Recent work discovered that a large fraction of mammary tumors from KB1P (*K14cre;Trp53^F/F^;Brca1^F/F^*) mice with acquired PARP inhibitor (PARPi) resistance featured downregulated expression of the non-essential histone variant H2AX^29,30^. Unexpectedly, subsequent elucidation of the underlying molecular mechanism of PARPi resistance in this model revealed strongly enhanced association of CtIP at stalled forks, ultimately restoring fork integrity in absence of functional BRCA1^29^. Therefore, we reasoned that *Brca1*-deficient KB1P tumor cells represent an interesting system to further substantiate our results. As shown previously^29^, SIRF analysis revealed significantly higher levels of spontaneous and HU-induced CtIP PLA foci in H2AX-depleted KB1P cells compared to cells transduced with a non-targeting (NT) gRNA (Figure 5A). Remarkably, pre-treatment with either PIN1 or p38α inhibitor, strongly abrogated CtIP association with stalled forks in both KB1P-derived cell lines (Figure 5A), substantiating the crucial role of PIN1 and p38α in CtIP isomerization to promote efficient localization of CtIP to sites of stalled forks.

**Figure 5.**
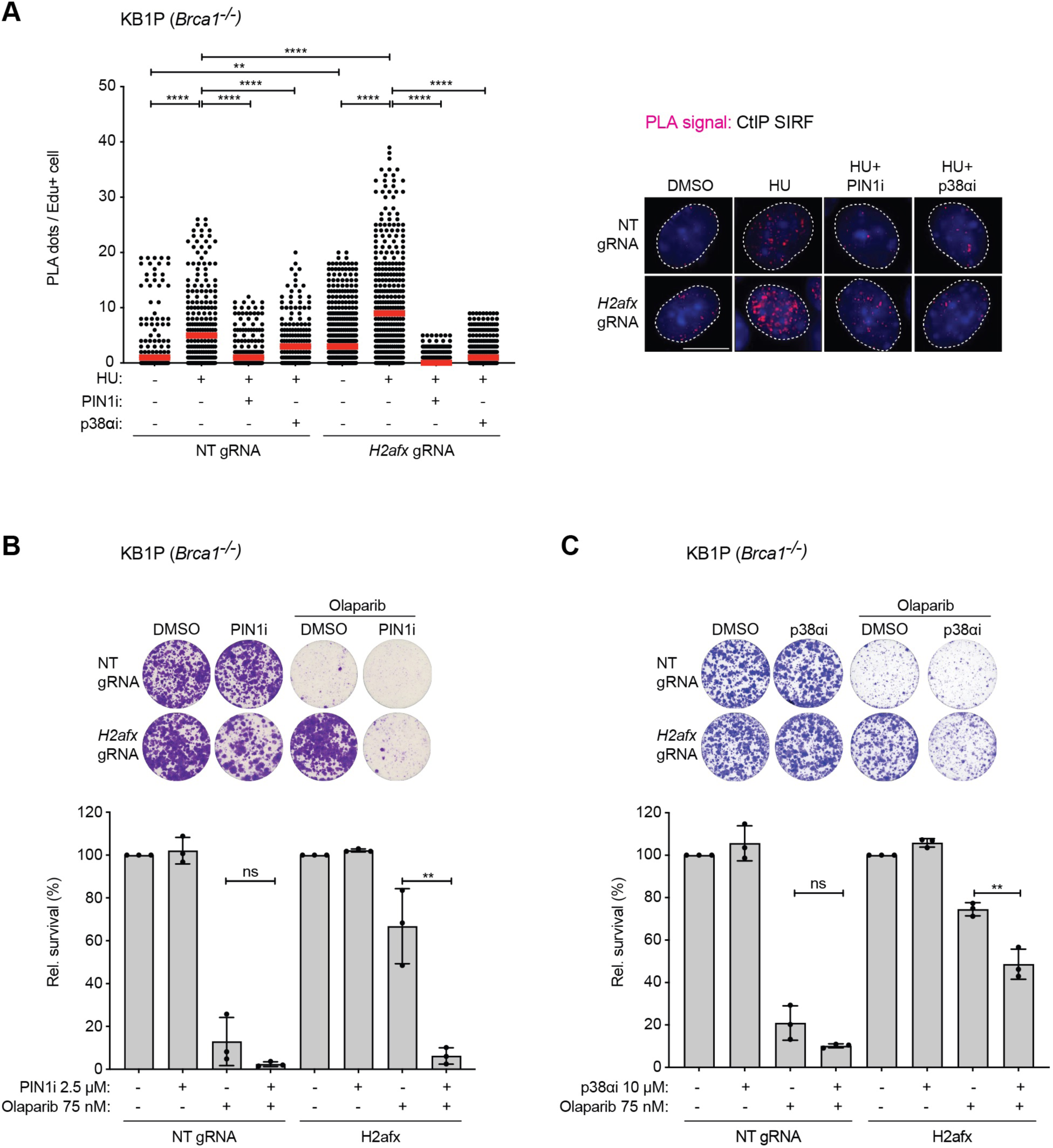
PIN1 or p38α inhibition impairs CtIP accumulation at stalled forks and overcomes Olaparib resistance in *Brca1^-/-^* tumor cells. **(A)** CtIP SIRF assay in KB1P-derived *Trp53^-/-^*; *Brca1^-/-^* and *Trp53^-/-^*; *Brca1^-/-^*; *H2afx^-/-^* cells, either mock-treated or treated with the PIN1 inhibitor KPT-6566 (10 µM) for 1h, or with the p38α inhibitor PH-797804 (1 µM) for 24h. Cells were pulse-labelled with EdU (25 µM) for 10 min followed by treatment with HU (8 mM) alone or in combination with the PIN1 or p38α inhibitors for 6h. Dot plots show the number of PLA foci and the median from at least 150 EdU-positive cells. Representative images are shown on the right. Scale bars, 10 μm. **(B)** Colony formation assay was performed in same cells as in (A), either mock-treated or treated with the PIN1 inhibitor KPT-6566 (2.5 μM) and with the PARP inhibitor Olaparib (75 nM) for 10 days. **(C)** Colony formation assay was performed in same cells as in (A), either mock-treated or treated with the p38α inhibitor PH-797804 (10 μM) and with the PARP inhibitor Olaparib (75 nM) for 10 days. **(B and C)** Plotted values are mean ± standard deviation of three biological replicates. Representative images are shown (top).

The development of PARPi resistance poses a great clinical challenge for the treatment of BRCA1/2-deficient tumors^31^. In recent years, several distinct mechanisms underlying PARPi resistance have been identified, including restoration of fork protection, providing new therapeutic strategies to potentially overcome PARPi resistance. Based on our findings, we therefore speculated whether the use of PIN1 or p38α inhibitors might represent such an opportunity. Strikingly, we observed that treatment with PIN1i significantly restored sensitivity to the PARPi Olaparib in H2AX-depleted KB1P cells (Figure 5B). As shown previously, we found that H2AX-deficient KB1P cells exhibit increased cellular resistance to chronic HU treatment (Figure S5A)^29^. Also here, PIN1 inhibition rendered BRCA1- and H2AX-deficient cells HU sensitive (Figure S5A). These findings reveal an unprecedented BRCA1- and HR-independent role of PIN1 in promoting chemoresistance, most likely through mediating CtIP-dependent restoration of fork protection in this context.

Finally, and consistent with an important role of p38α in promoting CtIP-dependent fork protection, combined treatment with the p38α inhibitor PH-797804 significantly restored both Olaparib and HU sensitivity in H2AX-deficient KB1P cells (Figures 5C and S5B). Collectively, these data corroborate the important role of PIN1-p38α signaling in promoting CtIP association with stalled forks and confirm a BRCA1-independent role of CtIP in fork stabilization that is governed by CtIP isomerization.

## DISCUSSION

We previously reported a distinct role for CtIP in the replication stress response by protecting regressed nascent DNA arms at forks from excessive digestion by the DNA2 nuclease^4^. While we could demonstrate that CtIP’s function in promoting DSB resection and HR repair is dispensable for fork stabilization, a broad understanding of the molecular mechanisms regulating CtIP-mediated fork protection remained to be established. Here, we reveal a critical role for phosphorylation-dependent prolyl isomerization of CtIP to protect HU-stalled forks from deleterious degradation. Based on our previous and present results, we propose a model in which CDK-mediated phosphorylation of CtIP-T315 during unperturbed S-phase enables PIN1 to recognize CtIP through its WW-phospho-binding domain. Subsequently, when cells are exposed to acute replication stress, p38α kinase phosphorylates CtIP at S276 to facilitate PIN1-mediated *cis*-to-*trans* isomerization of the S276-P277 peptide bond. This conformational change is required for efficient CtIP enrichment near stalled replication forks, which ultimately leads to the protection of nascent DNA at reversed forks from nucleolytic attack by DNA2 (Figure 6). Altogether, our data highlight CtIP isomerization as a molecular switch activating CtIP’s function in protecting reversed forks from nucleolytic degradation without compromising the resection and HR repair of DSBs.

**Figure 6.**
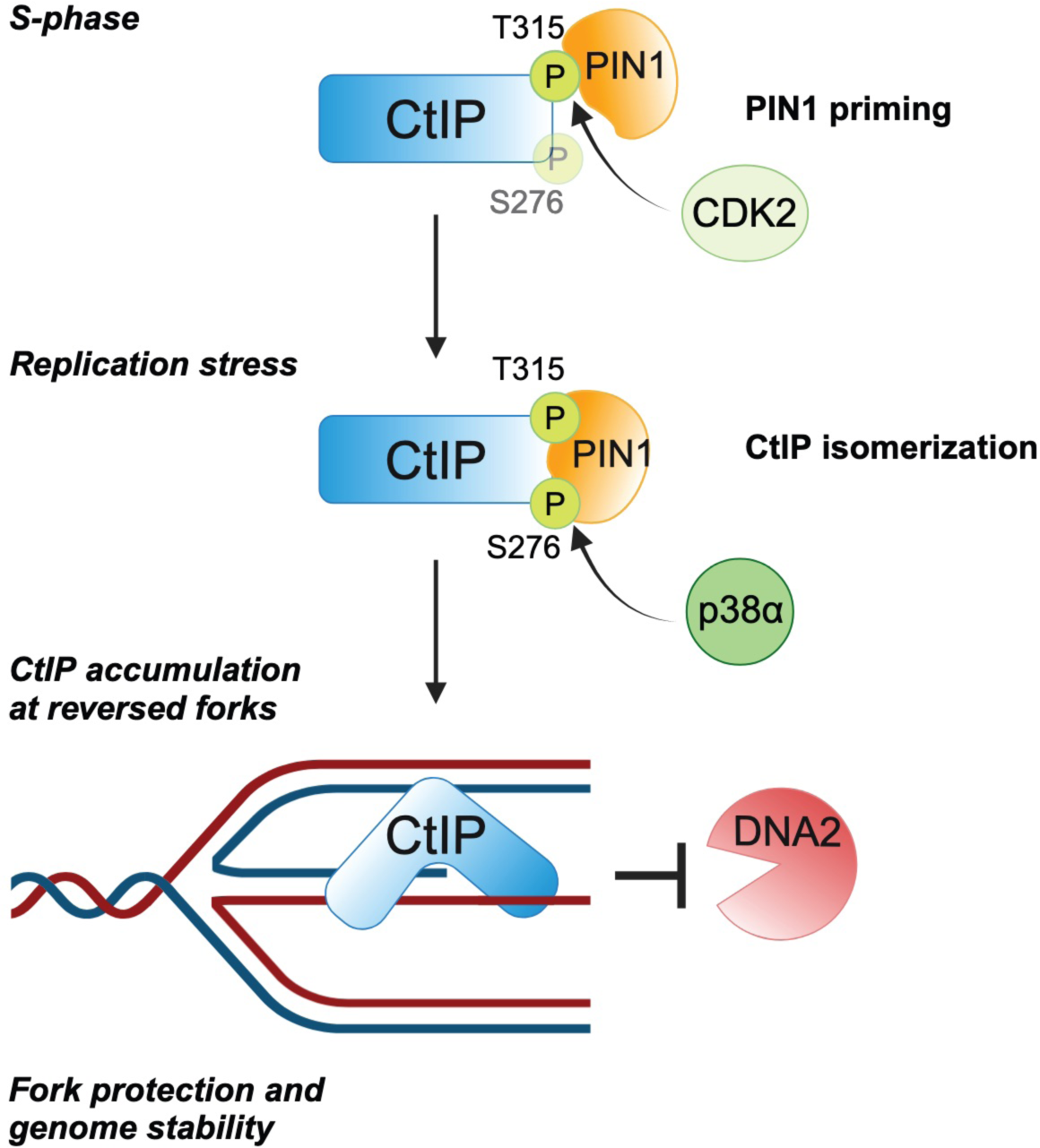
Schematic model depicting the role of PIN1-p38α-CtIP signaling in fork protection. During unperturbed S-phase, CDK2-mediated phosphorylation of T315 promotes PIN1 binding to CtIP. In response to replication stress, p38α kinase phosphorylates CtIP at S276. Subsequently, PIN1 catalyzes the *cis*-to-*trans* isomerization of the pS276-P277 peptide bond, ensuring accumulation of CtIP at stalled forks. Ultimately, this phosphorylation-isomerization cascade promotes CtIP-dependent protection of nascent DNA from DNA2-mediated nucleolytic processing, thereby maintaining of genome stability.

### Phosphorylation-dependent CtIP isomerization triggered by PIN1 and p38α promotes fork integrity

Earlier work from the Morris group demonstrated that fork protection by the BRCA1-BARD1 complex relies on PIN1-mediated isomerization of the BRCA1 pS114-P115 motif, which aids RAD51 recruitment to stalled forks, limiting nucleolytic processing of nascent DNA by MRE11^16^. Consistent with this work, we previously demonstrated that CtIP and BRCA1 act in separate fork protection pathways and synergistically alleviate replication stress-induced genomic instability, by restraining DNA2 and MRE11 fork resection activities, respectively^4^. We now provide evidence that, although counteracting fork degradation via two biochemically separable mechanisms, PIN1-mediated isomerization acts as a common upstream regulatory component controlling both CtIP- and BRCA1-dependent fork protection functions. Accordingly, we found that fork degradation induced by chemical inhibition of PIN1 isomerase activity is significantly, albeit not fully, rescued by the expression of a CtIP *trans*-locked mutant, indicating that BRCA1-mediated fork protection may be concomitantly compromised in this context.

Our previous work indicated that CDK-mediated T315 phosphorylation is a prerequisite for PIN1 recognition, while S276 phosphorylation is required for CtIP isomerization at the pS276-P277 site and mediated by a hitherto unknown proline-directed kinase. The MAPK family member p38α was reported to be activated in response to various sources of cellular stress, ranging from physiological situations (e.g. cell differentiation) to a wide range of exogenous and endogenous triggers, such as hyperosmolarity, oxidative stress or DNA damage^32^. For example, after exposure to ultraviolet (UV) radiation, p38α was reported to collaborate with CHK1 in the activation of the S-M checkpoint to prevent premature mitotic entry before completion of DNA replication^33^. This task was shown to gain even greater importance in p53-deficient cells, where ATM/ATR-dependent activation of p38α secures cell cycle arrest and cell survival in response to cisplatin, doxorubicin or camptothecin exposure^12^. Here, we uncover an unprecedented role for p38α in counteracting the degradation of HU-stalled forks through phosphorylating CtIP at S276, a requirement for subsequent *cis*-to-*trans* isomerization of the pS276-P277 prolyl peptide bond. Despite the presence of CtIP-pS276 in untreated conditions, we observed a further increase in S276 phosphorylation after HU exposure, which returned to control levels upon concomitant p38α inactivation, indicating that replication stress-induced CtIP-S276 phosphorylation relies on p38α. Importantly, our experiments were performed in non-synchronized U2OS cells, whereas robust activation of p38α was previously observed only when DNA-damaging agents were added in S-phase^23^. Strikingly, expression of the CtIP-S276A/P277A trans-locked variant completely restored fork stability in p38α-depleted cells, underscoring p38α-mediated S276 phosphorylation as a critical event in fork protection.

### Inhibition of the PIN1-p38α signaling axis restores chemosensitivity in *Brca1*-deficient mammary tumor cells

We previously demonstrated that combined depletion of CtIP and BRCA1 in U2OS cells provokes elevated levels of chromosomal instability, which are most likely attributed to the persistence of replication-associated DNA lesions^4^. Furthermore, reduced survival of BRCA1-deficient but not BRCA1-proficient cancer cells upon treatment with a CtIP-stapled peptide inhibitor suggested a synthetic sick relationship between BRCA1 and CtIP^34^. Recent work from Dibitetto and colleagues revealed that H2AX loss restores replication fork protection in *Brca1*-deficient mammary tumor cells via CtIP hyperaccumulation at stalled forks, resulting in PARPi resistance^29^. Consequently, CtIP inhibition using stapled peptides provoked fork degradation and restored chemosensitivity^29^. We now provide evidence that CtIP enrichment at HU-stalled forks in *Brca1*-deficient mouse tumor cells is compromised by PIN1 or p38α inhibition, indicating a pivotal role for the PIN1-p38α-CtIP signaling cascade as a critical regulator of fork stability in cells lacking functional BRCA1.

PIN1 ablation was previously reported to sensitize BRCA1-proficient breast cancer to PARPi as a result of impaired HR^19^. Here, we reveal that PIN1 and p38α inhibition restored Olaparib and HU sensitivity of *Brca1^-/-^;H2afx^-/-^* tumor cells that have acquired chemoresistance via restoration of fork protection but are still defective in HR. We observed a more pronounced effect of PIN1 inhibition in restoration of chemosensitivity compared to p38α inhibition, suggesting that besides BRCA1-BARD1 and CtIP, PIN1 is likely to engage more substrates than p38α implicated in fork protection. An interesting factor could be PTIP, which we identified in our previous proteomics analysis as a potential PIN1 interactor^11^. Work from the Nussenzweig group has shown that PTIP accumulates at sites of replication stalling and deposits MRE11 on stalled forks^35^. Accordingly, they found that PTIP loss promotes fork stability and chemoresistance in BRCA1-deficient cells though inhibition of MRE11-dependent fork degradation^35^. Taken together, PTIP’s function in response to replication stress might be negatively affected by isomerization and, thus, PIN1 inhibition could result in upregulated PTIP activity, resulting in MRE11 hyperaccumulation at stalled forks.

Finally, our findings also highlight that targeting the PIN1-p38α-CtIP axis might represent a promising therapeutic approach for BRCA1-mutated cancer that acquired chemoresistance. This strategy could also be relevant for pancreatic adenocarcinoma (PDAC), where PIN1 and p38 overexpression, as well as CtIP gene amplification, are frequently observed and found to correlate with poor prognosis^36–42^. Given the high prevalence of *KRAS* gain-of-function mutations in PDAC patients^43^, which endows those cancer cells with the ability to tolerate high levels of DNA damage and replication stress, we reason that targeting the PIN1-p38α-CtIP axis in pancreatic cancer may facilitate the development of improved therapies.

## METHODS

### Cell culture

U2OS and HEK293T, MEF and MEF *Pin1^-/-^* cells were grown in Dulbecco’s modified Eagle’s medium (DMEM, Thermo Fisher Scientific) supplemented with 10% fetal calf serum (FCS; GIBCO/Thermo Fisher Scientific) and 1% penicillin-streptomycin (Sigma-Aldrich). MDA-MB-436 cells were maintained in RPMI medium supplemented with 10% FCS and 1% penicillin-streptomycin. Cell lines were grown at 37° C in a humidified atmosphere with 6% CO_2_.

KB1P-G3 (*Trp53 -/-; Brca1 -/-* and *Trp53 -/-;Brca1 -/-;H2afx -/-*) cells were derived from KB1P mammary tumors as previously described^29^ and were grown at 37° C in 3% O_2_ in DMEM Nutrient mixture F-12 (Thermo Fisher Scientific) supplemented with 10% FCS, 1% penicillin-streptomycin, 5 ng/ml cholera toxin (Sigma-Aldrich), 5 μg/ml insulin (Sigma-Aldrich) and 5 ng/ml murine Epidermal Growth Factor (mEGF, Sigma-Aldrich). U2OS stably expressing GFP-empty vector (ev), GFP-CtIP-wt, S276A, T315A and S276A/T315A were generated as previously described^5^. U2OS cells inducibly expressing GFP-CtIP wt, S276A, P277A and S276A/P277A were generated as described below.

### Generation of U2OS GFP-CtIP doxycycline inducible cell lines

The Flp-In T-Rex system was used to generate U2OS cell lines stably expressing different siRNA-resistant GFP-CtIP constructs under the control of doxycycline-inducible promoter like described before^44^. In brief, U2OS Flp-In T-Rex cells were transfected with expression vectors pcDNA5/FRT/TO-GFP-CtIP wt, S276A, P277A and S276A/P277A and the Flp recombinase expression plasmid, pOG44, mixed in a 1:9 ratio using FuGENE6 Transfection Reagent. Cells were plated 24 h post transfection. The next day, the medium was supplemented with 250 μg/mL hygromycin B (InvivoGen) and 15.5 μg/mL blasticidin S (InvivoGen). GFP-positive bulk cultures were sorted using a BD FACSAria III cell sorter (Flow Cytometry Facility, University of Zurich). Sorted cell lines were tested for expression and nuclear localization of the transgene-products via immunofluorescence microscopy and western blotting analysis. Induction of GFP-CtIP expression was performed by growing the inducible cell lines for 24 h in medium supplemented with 1 μg/ml of doxycycline.

### Generation of U2OS GFP-CtIP hypomorphic cells

U2OS^Cas9/*CtIP*^ cell lines stably expressing GFP-tagged wt and mutant CtIP were generated as previously described^4^. Briefly, U2OS^Cas9/*CtIP*^ cells were transfected with the appropriate plasmids and with a puromycin resistant cassette containing pcDNA5/TO vector in a 5:1 ratio. Selection of GFP-positive cells was performed complementing the medium with 1 μg/mL puromycin (InvivoGen/LabForce). GFP-positive bulk cultures were tested for expression and nuclear localization of the transgene-products by immunofluorescence microscopy and western blotting analysis.

### Plasmids and cloning

DNA primers used for cloning and sequencing were obtained from Microsynth (Balgach, Switzerland). pEGFP-C1 plasmids containing CtIP wt and S327A were previously described^45^. The pEGFP-C1 plasmid containing CtIP-S276A, P277A and S276A/P277A were generated by site-directed mutagenesis. pcDNA5/FRT/TO-GFP expressing CtIP-wt has been previously described^44^. The S276A, P277A and S276A/P277A mutants of CtIP in pcDNA5/FRT/TO-GFP vector were generated by site-directed mutagenesis. The FLAG-CtIP wt and T847A expression vectors were described previously^44,46^. The S276A, P277A and S276A/P277A mutants of CtIP in the FLAG vector were generated by site-directed mutagenesis. All constructs were verified by sequencing. Primers used for site-directed mutagenesis are reported below with 5’ to 3’ orientation:

CtIP_S276A _Forward: AAGGTCCCATGGCCCCCCTTGGTGATGAGCTCTAC

CTIP_S276A _Reverse: CACCAAGGGGGGCCATGGGACCTTGAGTTTCAGA

CtIP_P277A_Forward: CATGAGCGCTCTTGGTGATGAGCTCTACCACTGTC

CtIP_P277A_Reverse: CCAAGAGCGCTCATGGGACCTTGAGTTTCAG

CtIP_S276A/P277A_Forward: CCATGGCTGCCCTTGGTGATGAGCTCTACCAC

CtIP_S276A/P277A_Reverse: CAAGGGCAGCCATGGGACCTTGAGTTTCAG

### siRNA transfections and sequences

siRNA oligos were transfected using Lipofectamine RNAiMAX (Invitrogen) according to manufacturer’s instructions, at a final concentration of 10 nM or 40 nM. Experiments were performed 48 h post siRNA transfection.

### Drug treatments

The following compounds were used at the indicated final concentrations: hydroxyurea (HU, 80 µM in colony formation assay, 2 mM and 8 mM in DNA fibers and SIRF, 5 mM in metaphase spreads), cycloheximide (100 µg/mL), mirin (25 µM in DNA fibers), DNA2i NSC-105808 (2 µM in DNA fibers), PIN1i KPT-6566 (2.5 and 7.5 µM in colony formation assay and 5 and 10 µM in DNA fibers, HR assays and laser micro-irradiation), p38αi PH-797804 (1 µM in DNA fibers, SIRF and immunoprecipitation, 10 µM in colony formation assay), p38α Protac NR-11c (1 µM in DNA fibers and immunoprecipitation), Olaparib (75 nM in colony formation assay).

### Immunoblotting and triton extraction

For western blotting analysis, cell extracts were prepared in Laemmli buffer (4% SDS, 20% glycerol, 120 mM Tris-HCl pH 6.8). Chromatin-enriched lysates were performed as previously described^45^. In brief, cells were washed with cold PBS and incubated 5min at 4°C with pre-extraction buffer (25 mM HEPES pH 7.4, 50 mM NaCl, 1 mM EDTA, 3 mM MgCl2, 300 mM sucrose, 0.5% Triton X-100 and protease inhibitors). Adherent cellular material was collected in Laemmli buffer. After heat-denaturation of the chromatin enriched fraction, lysates were sonicated and analyzed by western blotting.

For immunoblotting proteins were resolved by SDS–PAGE and transferred to nitrocellulose membranes. Membranes were incubated at 4°C overnight with the appropriate primary antibodies and 1 h at room temperature (RT) with secondary antibodies. Proteins were then visualized with the Advansta WesternBright ECL reagent and the VilberLourmat Fusion Solo S imaging system.

### Immunoprecipitation

For immunoprecipitation (IP), GFP-Trap and Myc-Trap (ChromoTek, proteintech) cells were lysed in NP-40 extraction buffer [50 mM Tris-HCl, pH 7.5, 120 mM NaCl, 1 mM EDTA, 6 mM EGTA, 15 mM sodium pyrophosphate and 1 % NP-40 supplemented with phosphatase inhibitors (20 mM NaF, 1 mM sodium orthovanadate) and protease inhibitors (1 mM benzamidine and 0.1 mM PMSF, Protease inhibitor cocktail, Sigma-Aldrich)]. Cell lysates were incubated with Benzonase (Merck) for at least 30 min at 4 °C cleared by centrifugation and protein concentration was determined by Bradford assay (Bio-Rad).

1-2mg of cleared lysates were incubated with ChromoTek GFP/Myc-Trap Agarose beads (proteintech) for 2h and washed three times with NTEN300 buffer (0.5% NP-40, 0.1 mM EDTA, 20 mM Tris-HCl pH 7.4, 300 mM NaCl) or three times with NP-40 extraction buffer and once with TEN100 buffer (20 mM Tris-HCl pH 7.4, 0.1 mM EDTA and 100 mM NaCl).

For endogenous IPs lysates were incubated at 4°C overnight with 1 ug of antibody per milligram of lysates. Protein A beads (GE Healthcare) were added afterwards for 2 h and washed as described above.

Complexes bound to beads were boiled in SDS sample buffer and analyzed by SDS–PAGE followed by western blotting analysis as described above.

### Antibodies

For DNA fiber assay the following antibodies are used: mouse anti-BrdU/IdU 1:80, BD Biosciences 347580; Rat anti-BrdU/CldU 1:400, Abcam ab6326; goat anti-mouse Alexa Fluor™ 488 1:250, Thermo Fisher Scientific; donkey anti-rat Cy3 1:250, Jackson ImmunoResearch.

For SIRF the following antibodies are used: rabbit anti-CtIP 1:100, Bethyl Laboratories #A300-488A; mouse anti-Biotin 1:200, #200-002-211, Jackson Immuno Research; rabbit anti-Biotin 1:1000, A150-109A, Bethyl; mouse anti-GFP 1:100, Roche 11814460001 IgG1κ clones 7.1 and 13.1; goat anti-mouse Alexa Fluor™ 488 1:1000, Thermo Fisher Scientific #A11029 and goat anti-rabbit Alexa Fluor™ 546 1:1000, Thermo Fisher Scientific #A11010.

For QIBC the following antibodies are used: rabbit anti-RPA32 1:500, Abcam ab76420; donkey anti-rabbit Alexa Fluor™ 647 1:500, Thermo Fisher Scientific #A31573.

For immunoblotting the following antibodies are used: mouse anti-Myc (9E10) 1:500, Thermo Fisher Scientific MA1-980; mouse anti-CtIP (D4) 1:250, Santa Cruz sc-271339; rabbit anti-CtIP (D76F7) 1:1000, Cell Signaling #9201; rabbit anti-pS276-CtIP 1:200 custom made with Eurogentec with synthetic phosphopeptides (KLH-coupled) corresponding to residues surrounding S276 (ETQGPMpSPLGDEL)^11^; mouse anti-Mre11 1:1000 Genetex #GTX70212; mouse anti-p38a 1:1000 Cell Signaling #9217; rabbit anti-p38a 1:1000 Cell Signaling #9218; rabbit anti-Phospho-p38 MAPK T180/Y182 Cell Signaling #9211; mouse anti-FLAG M2 1:1000, Sigma-Aldrich F1804; rabbit anti-Lamin B1 1:1000 ab16048; mouse anti-Tubulin 1:20’000 Sigma-Aldrich #T9026; rabbit anti-GFP 1:1000, Abcam ab290; mouse anti-GFP (B2) 1:500, Santa Cruz sc-9996; rabbit anti-Cyclin D1 1:1000, Cell Signaling #2922; rabbit anti-SMARCAL1 1:1000, Abcam ab154226; mouse anti-GAPDH 1:40’000, Millipore MAB374; mouse anti-BRCA1 (D9) 1:50, Santa Cruz sc-6954.

### DNA fiber analysis

DNA fiber analyzes were performed as described previously^47,48^. In brief, non-synchronized U2OS cells were labeled with CldU (33 μM) for 30 min, followed by IdU (340 µM) for 30 min before incubation with HU for 4 h. Alternatively, cells were labeled with CldU for 20 min, subsequently treated with HU for 2 h and chased with IdU for 40 min before harvesting in PBS. Cells lysis was performed (lysis buffer: 200 mM Tris-HCl (pH 7.4), 50 mM EDTA, 0.5% SDS) and DNA fibers were stretched onto glass slides, air-dried at RT for 30 min and fixed in Methanol:Acetic acid in a 3:1 ratio (Merck) at 4°C overnight. Fibers were rehydrated in PBS before denaturation with 2.5 M HCl for 1 h, washed with PBS and blocked with 2% BSA in PBS+0.1% Tween 20 for 45 min. The CldU and IdU tracks were immunostained using anti-BrdU primary and corresponding secondary antibodies. Coverslips were mounted using ProLong Gold Antifade Mountant (Life Technologies). Images were acquired on a Leica DMI 6000 fluorescence microscope using 63x objective and analyzed using Fiji software^49^.

### Metaphase spreads

Metaphase spreads were performed as described previously^50^. Briefly, 0.1 µg/mL colcemid was added to the cells 2 h prior harvesting by trypsinization. Cell pellets were resuspended in 5 ml of hypotonic solution (potassium chloride 75 mM) and incubated at 37°C for 30 min for swelling. Cells were then fixed a first time for 3 min with 5% acetic acid and then two times for 10 min with ethanol-acetic acid in a 3:1 ratio. Fixed cells were gently resuspended in fixative solution to achieve optimal cell density before dropping onto glass slides. Slides were mounted using Vectashield® Mounting Media (Vector Laboratories) containing 4’,6-Diamidino-2-Phenylindole Dihydrochloride (DAPI). Fluorescent images were acquired using a Leica DMI 6000 fluorescence microscope with 63x objective.

### HR reporter assay

HR reporter assay was carried out as described previously^11,51^. In brief, U2OS EGFP-HR were seeded into 10 cm dishes and transfected with siRNA control (siCNTL) or targeting CtIP (siCtIP). AÄer 24 h, cells were seeded into 12-well plate. The next day, cells were transfected with pcDNA3 or *I-SceI* expression plasmid (pCBASce) and FLAG-ev (empty vector), FLAG-CtIP-wt, S276A, P277A, S276A/P277A and T847A using jetPrime transfection reagent (Polyplus). For the experiments shown in figure S2D and S2G, cells were directly seeded into 12-well plates and treated with PIN1i KPT-6566 3 h before transfection with pcDNA3 or *I-SceI* expression plasmid (pCBASce) and the indicated FLAG-CtIP constructs. For all experiments, medium was exchanged 4 h aÄer transfection and cells were harvested 48 h post-transfection. As read out for HR, GFP expression was measured by flow cytometry using AÇune Nxt Flow Cytometer equipped with a 488 nm laser and 530/30 band-pass filter. A minimum of 20’000 events per sample were recorded.

### Expression and purification of recombinant proteins

The CtIP-S276A variant was prepared by mutating the respective wild-type pFB-2xMBP-CtIP-10xhis plasmid by QuickChange site-directed mutagenesis kit following manufacturer’s instructions (Agilent Technology). The wild-type protein, as well as the point mutant, were expressed in Sf9 insect cells in SFX Insect serum-free medium (Hyclone) using the Bac-to-Bac expression system (Invitrogen), according to manufacturer’s recommendations. Purification was performed by affinity chromatography exploiting the N-terminal maltose-binding protein (MBP)-tag and the C-terminal 10xhis-tag^52^. For expression of phosphorylated CtIP (pCtIP) variants, Sf9 cells were treated with 50 nM Okadaic acid (APExBIO) to preserve proteins in their phosphorylated state, and 1 μM camptothecin (Sigma) to further activate protein phosphorylation cascade. The MRN complex was prepared using the 3xflag-tag at the C-terminus of RAD50.

### Preparation of oligonucleotide-based substrate

All oligonucleotides were purified by polyacrylamide gel electrophoresis and purchased from Eurogentec. The labeling of oligonucleotides at the 5’-end was carried out by T4 polynucleotide kinase (New England Biolabs) and [γ-32P] ATP (Hartmann Analytic). To prepare quadruple blocked 70-bp long DNA substrate, PC210 and PC211 oligonucleotides were used, as described previously^53^.

### Endonuclease assay with recombinant proteins

Endonuclease assays (15 μl volume) were performed in nuclease buffer containing 25 mM Tris-HCl pH 7.5, 5 mM magnesium acetate, 1 mM manganese acetate, 1 mM dithiothreitol (DTT), 1 mM ATP, 0.25 mg/ml BSA (New England Biolabs) and 1 nM oligonucleotide-based DNA substrate (in molecules). The reactions were supplemented with 15 nM monovalent streptavidin and incubated for 5 min at RT to block the biotinylated ends of the DNA substrates. The recombinant proteins were then added to the reactions on ice and samples were incubated at 37°C for 2 h. Reactions were stopped by adding 0.5 μl ethylenediaminetetraacetic (0.5 M EDTA) and 1 μl Proteinase K (19 mg/ml, Roche), and incubated at 50°C for 30 min. Finally, 16.5 μl loading buffer (5% formamide, 20 mM EDTA, bromophenol blue) was added to all samples and the products were separated on 15% polyacrylamide denaturing urea gels (19:1 acrylamide-bisacrylamide, Bio-Rad). The gels were fixed in fixing solution (40% methanol, 10% acetic acid, 5% glycerol) for 30 min at room temperature and dried on a 3MM Chr paper (Whatman). The dried gels were exposed to storage phosphor screen (GE Healthcare) and scanned by a Typhoon Phosphor Imager (FLA 9500, GE Healthcare).

### SIRF (in Situ analysis of protein Interactions at DNA Replication Forks)

SIRF assay was performed as previously reported^26,27^. Briefly, cells were seeded on coverslips and, after 24 h, pulsed-labelled with 25μM EdU for 10 min. Afterwards, cells were washed three times with PBS to remove the EdU and either incubated with HU and the indicated inhibitors or immediately pre-extracted and fixed (for untreated samples). Pre-extraction was performed with CSK buffer containing 0.5% of Triton™ X-100 (Sigma-Aldrich) on ice for 5 min and fixation was done with 4% Paraformaldehyde at RT for 15 min. Coverslips were then washed with PBS and stored overnight at 4°C. The following day, EdU was chemically linked to Biotin-azide using the Click-iT™ Reaction Kit (Thermo fisher scientific) for 1 h at 37°C. In situ proximity ligation assay (PLA) was performed using Duolink PLA technology (Sigma-Aldrich) according to the manufacture instructions. In brief, coverslips were blocked for 1h at 37°C with blocking solution, followed by incubation with primary antibodies for 2 h at RT. After primary antibody incubation, coverslips were washed with Wash Buffer A (0.01 M Tris, 0.15 M NaCl and 0.05% Tween 20) for 5 min at RT and incubated for 1 h at 37°C with Duolink anti-Mouse PLUS and anti-Rabbit MINUS PLA probes. After three wash steps in Wash Buffer A for 5 min, PLA probes were ligated for 30 min at 37°C. Coverslips were then washed three times 5 min in Wash Buffer A. Amplification was performed using the ‘Duolink In Situ Detection Reagents FarRed’ (Sigma-Aldrich) at 37°C for 100 min. After amplification, coverslips were washed twice in Wash Buffer B (0.2 M Tris and 0.1 M NaCl) for 10 min and incubated for 30 min at 37°C with the appropriate secondary antibody. Coverslips were then washed twice with Wash Buffer B and once in 0.01x Wash Buffer B for 1 min. Finally, coverslips were mounted using Vectashield® Mounting Media (Vector Laboratories) containing DAPI, sealed and imaged on a Leica DMI 6000 fluorescence microscope using a 63x objective. Analysis of PLA foci in EdU positive cells was performed using CellProfiler.

### Laser micro-irradiation coupled live cell imaging

Cells were seeded on a glass-bottom chambered coverslip (Ibidi), treated with 10 µM 5-bromo-2’-deoxyuridine for 24 h. Samples were imaged on an inverted confocal spinning disk microscope [Olympus IX83] equipped with CSU-W1 unit [Yokogawa, Japan] SoRa disk for super resolution imaging, using a 60X [Olympus, Japan] objective, under controlled temperature (37°C) and CO_2_ (5%) (Cellvivo incubation system). Additionally, cells were irradiated with a pulsed 355 nm laser [UGA 42 Caliburn, Rapp OptoElectronic, Wedel, Germany]. Cells were imaged with a 488nm laser, the emission wavelength range was 500-550 nm (BP 525/50). Time-lapse images were capture for the indicated time intervals.

The media used during the live imaging is the Gibco™ FluoroBrite™ DMEM complemented with 10% FCS (GIBCO) and 1% penicillin-streptomycin (Sigma-Aldrich). The signal intensity of the irradiated path was calculated using ImageJ software.

### High-content microscopy and quantitative image-based cytometry (QIBC)

U2OS and U2OS^Cas9/*CtIP*^ cell lines stably expressing GFP-tagged WT and mutant CtIP or transfected with the indicated siRNAs were grown on sterile 12mm glass cover slips. Typically, after indicated treatment or siRNA transfection, cells were then fixed in 3% formaldehyde for 15 min at room temperature, washed once in PBS, permeabilized for 5 min at room temperature in 0.2% Triton™ X-100 (Sigma-Aldrich) in PBS, washed twice in PBS and incubated in blocking solution (filtered DMEM containing 10%FBS and 0.02% Sodium Azide) for 15 min at room temperature. To detect chromatin-associated RPA2 levels, cells were pre-extracted in 0.2% Triton™ X-100 in PBS for two min on ice prior to formaldehyde fixation. For antibody staining, cells were incubated in blocking solution with primary antibodies for 2 h at room temperature, washed three times with PBS and incubated with secondary antibodies in blocking solution for 1h at room temperature. Cells were washed once with PBS and incubated for 10 min with DAPI (0.5 mg/ml) in PBS at room temperature. Following three washing steps in PBS, coverslips were briefly washed with distilled water and mounted on 5 ml Mowiol-based mounting media [Mowiol 4.88 (Calbiochem) in Glycerol/TRIS)]. Automated multichannel wide-field microscopy for high-content imaging and quantitative image-based cytometry (QIBC) was performed using the Olympus ScanR System as described previously^54^. Images were analyzed with the inbuilt Olympus ScanR Image Analysis Software Version 3.3.0, a dynamic background correction was applied, and nuclei segmentation was performed using an integrated intensity-based object detection module based on the DAPI signal. All downstream analyzes were focused on properly detected nuclei containing a 2C-4C DNA content as measured by total and mean DAPI intensities. Fluorescence intensities were quantified and are depicted as arbitrary units. Color-coded scatterplots of asynchronous cell populations were generated with Spotfire data visualization software (TIBCO Spotfire 10.10.1.7). Within one experiment, similar cell numbers were compared for the different conditions. For visualizing discrete data in scatterplots, mild jittering (random displacement of data points along discrete data axes) was applied to demerge overlapping data points. Representative scatterplots and quantifications of independent experiments, typically containing several thousand cells each, are shown.

### Colony formation assay

KB1P cells were seeded in 6-well plates at 4,000 cells/well to assess survival upon treatment with Olaparib or HU. Cells were either mock treated (DMSO) or treated with the indicated concentrations of Olaparib, HU, PIN1 inhibitor KPT-6566 or p38α inhibitor PH-797804 the day of seeding. The treatment lasted for the whole duration of the experiment and was refreshed twice a week. After 10 days of growth, cells were fixed with crystal violet solution [0.5% crystal violet and 20% ethanol (w/v)]. Plates were scanned and survival was analyzed with the ImageJ plugin Colony Area using the parameter colony intensity as readout.

### Quantification and Statistical Analysis

For QIBC analysis a total of 20 images with 20x objective were acquired in an unbiased fashion from asynchronous cell population. Typically, between 1000 and 3000 cells per condition were analyzed, and representative single cell data of cell cohorts of comparable size are shown as one-dimensional cell cycle-resolved scatterplots. Fluorescence intensities were quantified and are depicted as arbitrary units. Color-coded scatterplots of asynchronous cell populations were generated with Spotfire data visualization software (TIBCO). Within one experiment, similar cell numbers were compared for the different conditions. For visualizing discrete data in scatterplots, mild jittering (random displacement of data points along discrete data axes) was applied in order to demerge overlapping data points. Representative scatterplots and quantifications of independent experiments are shown.

Statistical analyses were performed using GraphPad Prism (GraphPad Software Inc). For HR assay and colony formation assay p values were calculated with the unpaired t-test. When comparing more than two groups, one-way ANOVA was used. For DNA fibers experiments a minimum of 110 fibers were scored per sample. Each experiment was repeated at least twice, and representative experiments are shown. The samples were subjected to a Mann-Whitney analysis. In all cases: ****P ≤ 0.0001; ***P ≤ 0.001; **P ≤ 0.01; *P ≤ 0.05, ns, non-significant.

## ACKNOWLEDGEMENTS

We wish to thank Sara Przetocka and all members of the Sartori lab for critical reading of the manuscript. We thank MaÇhias Altmeyer, Richard Chahwan and Massimo Lopes for helpful discussions. We thank Daniel Gonzalez Acosta for technical assistance with establishing the SIRF assay. We thank Giannino Del Sal for providing *Pin1^-/-^* MEFs. We are grateful to the Flow Cytometry Facility and the Center for Microscopy and Image Analysis at the University of Zurich for the sorting of U2OS cell lines and for technical support, respectively. Financial support came from the Swiss National Science Foundation (31003A_176161 and 310030_208143 to A.A.S., 320030M_219453 to S.R., 310030_207588 and 310030_205199 to P.C.), the European Union (ERC-2019-AdG-883877 to S.R., ERC-2020-AdG-101018257 to P.C.), the UZH Candoc Grant (no. FK-23-050 to F.V.) and the AIRC Fellowship for Abroad (to L.M.). Figure 6 was created with BioRender.

## AUTHOR CONTRIBUTIONS

Conceptualization: F.V. and A.A.S.

Investigation: F.V., M.G., L.M., H.D., A.P., G.C., I.C., C.v.A., V.v.A., S.W., B.C., M.C-R. and D.D.

Resources: M.S., A.R., A.R.N., P.C. and S.R.

Writing – Original Draft: F.V. and A.A.S.

Writing – Review & Editing: F.V., M.G., A.R.N. and A.A.S.

Supervision, Project Administration & Funding Acquisition: A.A.S.

## SUPPLEMENTAL INFORMATION LEGENDS

**Figure S1.**
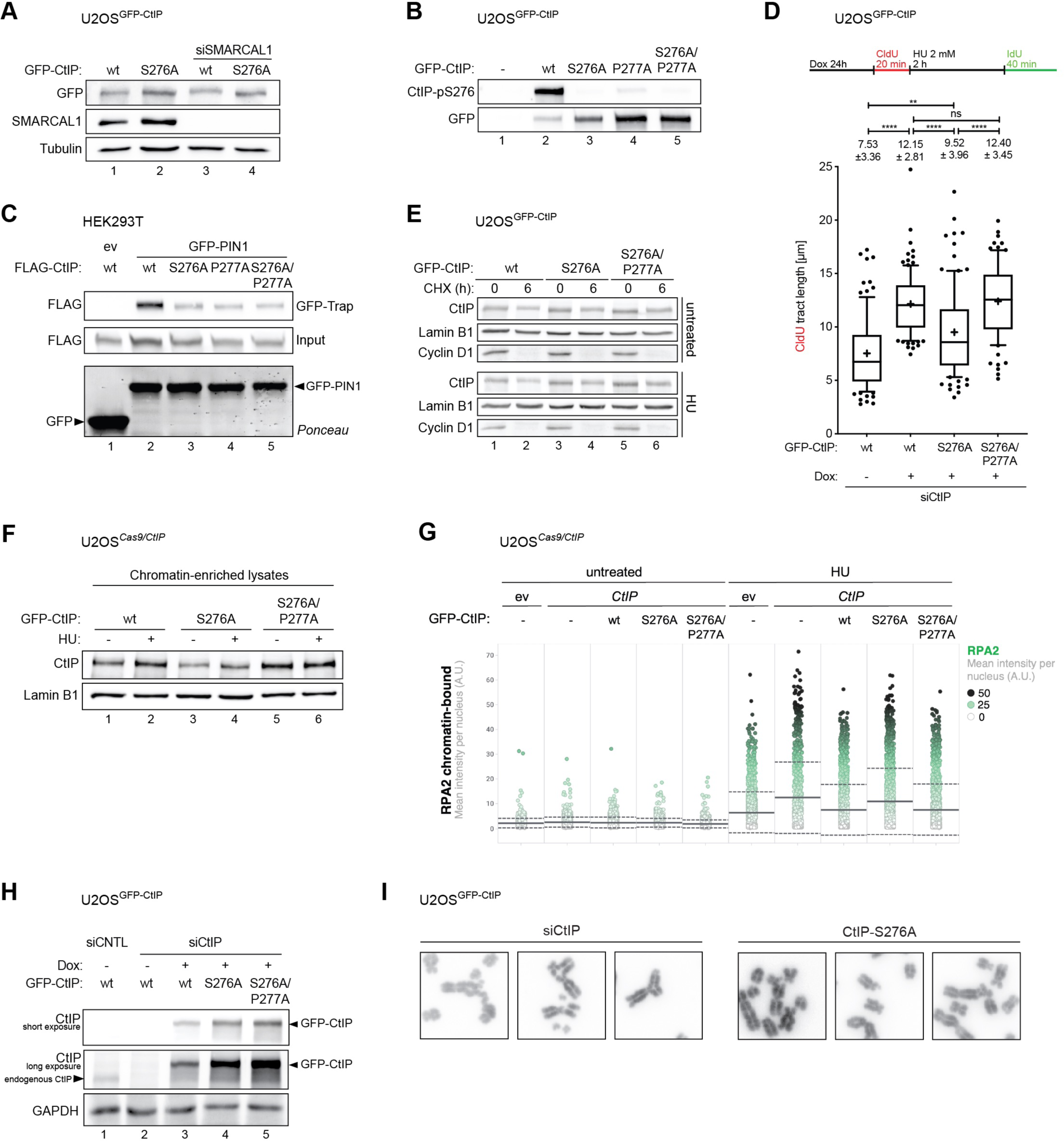
**(Related to Figure 1). (A)** Western blotting of lysates from U2OS cells inducibly expressing GFP-CtIP wt or S276A and depleted of endogenous CtIP alone, or co-depleted of CtIP and SMARCAL1. **(B)** GFP-Trap of U2OS cells inducibly expressing GFP-CtIP variants and depleted of endogenous CtIP. Whole-cell lysates (input) and immunoprecipitates were analyzed by western blotting using specific antibodies. **(C)** GFP-Trap of HEK293T cells co-transfected with GFP-PIN1 and indicated FLAG-CtIP variants. Whole-cell lysates (input) and immunoprecipitates were analyzed by western blotting using specific antibodies. **(D)** Fork degradation was evaluated upon HU treatment in U2OS cells depleted of endogenous CtIP and stably expressing indicated GFP-CtIP variants. Box and whisker plots of CldU-tract length for individual replication forks are shown. Numbers indicated above the individual plots represent the mean tract length ± standard deviation. Schematics of the CldU/IdU pulse-labelling protocol are shown (top). **(E)** Western blotting of lysates from U2OS cells inducibly expressing GFP-CtIP wt, S276A and S276A/P277A *trans*-locked mutant were either mock-treated or treated with HU (2 mM) for 4h. Cells were then released into fresh medium supplemented with cycloheximide (CHX, 100 µg/ml) for 6h, and lysates were analyzed by western blotting with the indicated antibodies. **(F)** Western blotting of chromatin-enriched lysates of U2OS*^Cas9/ev^*and U2OS*^Cas9/CtIP^* cells complemented with indicated GFP-CtIP variants. **(G)** Quantitative image-based cytometry (QIBC) of chromatin-loaded RPA2 in U2OS*^Cas9/ev^* and U2OS*^Cas9/CtIP^*cells complemented with indicated GFP-CtIP variants and treated or not with HU (2 mM for2h). Chromatin-bound RPA2 mean intensities are plotted and color-coded. The mean (solid line) and standard deviation (dashed line) are indicated. n > 1’500 cells per condition from minimum of two biological replicates. **(H)** Western blotting of lysates from U2OS cells inducibly expressing indicated GFP-CtIP variants and depleted of endogenous CtIP as employed in the metaphase spread analysis. **(I)** Representative images of HU-induced chromosomal aberrations typically observed in U2OS cells transfected with siCtIP or of CtIP-depleted cells expressing siRNA-resistant CtIP-S276A.

**Figure S2.**
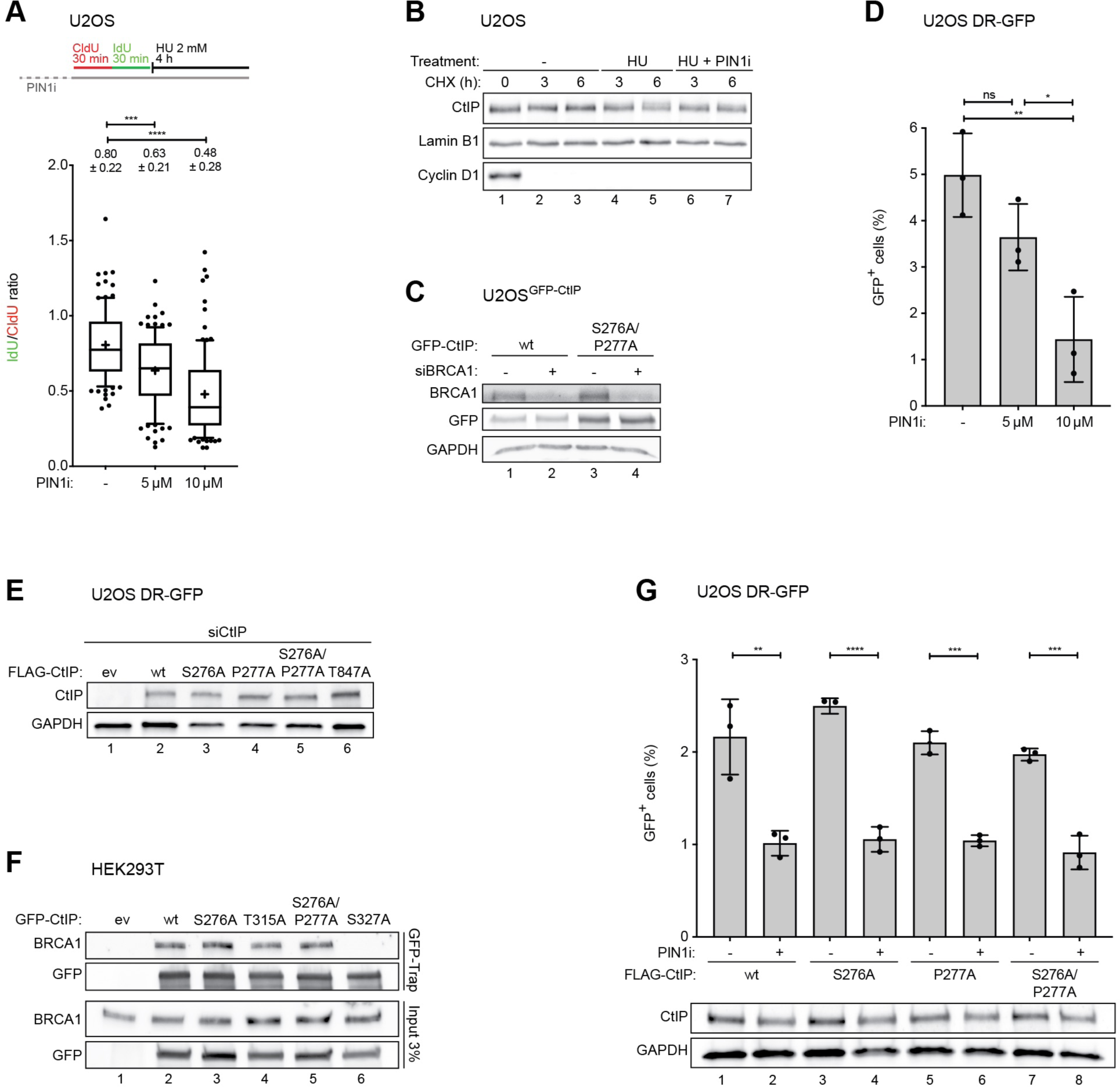
**(Related to Figure 2). (A)** Fork degradation was evaluated upon HU treatment in U2OS cells pre-treated for 1h with the PIN1 inhibitor KPT-6566. Box and whisker plots of IdU/CldU-tract length ratios for individual replication forks are shown. Numbers indicated above the individual plots represent the mean ratios ± standard deviation. Schematics of the CldU/IdU pulse-labelling protocol are shown (top). **(B)** Western blotting of lysates from U2OS cells were either mock-treated, treated with HU (2mM, 4h) or with HU and PIN1 inhibitor (10µM, 1h before HU treatment). Cells were then released into fresh medium supplemented with cycloheximide (CHX, 100 µg/ml) for the indicated time points and analyzed using specific antibodies. **(C)** Western blotting of lysates from U2OS cells inducibly expressing indicated GFP-CtIP variants and depleted of endogenous CtIP alone or in combination with BRCA1 depletion and analyzed using specific antibodies. **(D)** HR efficiency was evaluated in U2OS/DR-GFP cells mock-treated or treated for 3h with the indicated concentrations of the PIN1 inhibitor KPT-6566 before transfection with the *I-SceI* expression plasmid. Cells were harvested at 48h post-transfection and analyzed by flow cytometry for GFP signal. Data are shown as percentage of GFP-positive cells. **(E)** Western blotting of lysates from U2OS/DR-GFP cells depleted for endogenous CtIP and transfected with either empty vector (ev) or indicated FLAG-CtIP constructs. **(F)** GFP-Trap of HEK293T cells transfected with indicated GFP-CtIP variants. Whole-cell lysates (input) and immunoprecipitates were analyzed by western blotting using specific antibodies. **(G)** HR efficiency was evaluated in U2OS/DR-GFP cells mock-treated or treated with the PIN1 inhibitor KPT-6566 3h before co-transfection with the *I-SceI* expression plasmid and indicated FLAG-CtIP constructs. Cells were harvested at 48h post-transfection and analyzed by flow cytometry for GFP signal. Data are shown as percentage of GFP-positive cells. Western blotting of lysates from the same experiment is shown below.

**Figure S3.**
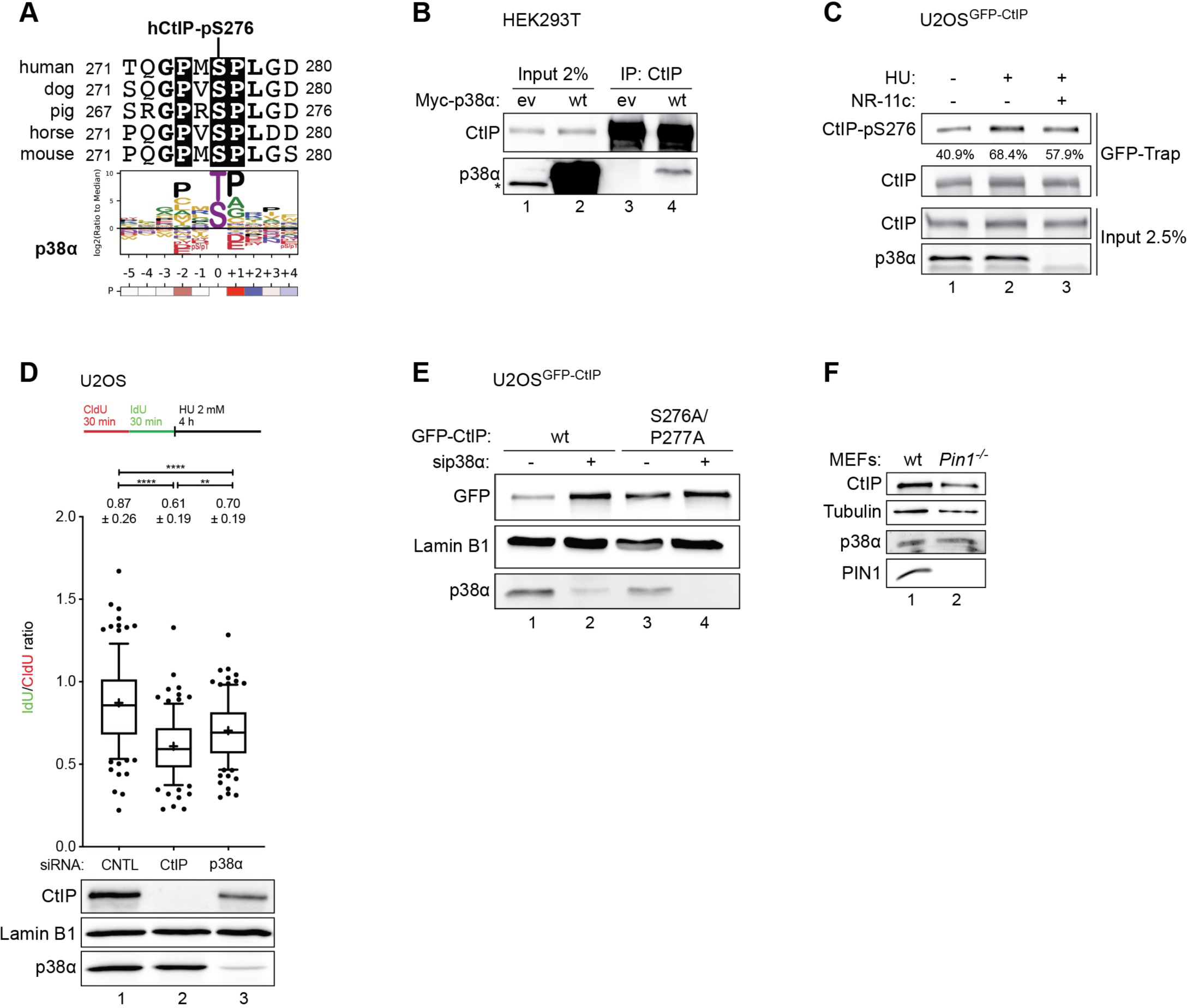
**(Related to Figure 3). (A)** Multiple sequence alignment of the CtIP region containing S276. The full consensus sequence for p38α substrates is shown below (modified from Johnson et al., 2023^55^). **(B)** Immunoprecipitation of endogenous CtIP from HEK293T cells transfected with Myc-p38α. Whole-cell lysates (input) and immunoprecipitates were analyzed by western blotting using specific antibodies. The * indicates an unspecific band. **(C)** GFP-Trap of U2OS cells inducibly expressing GFP-CtIP and treated with HU (2 mM, 4h). Where indicated, cells were treated with the p38α PROTAC NR-11c (1 µM, 24h before HU). Whole-cell lysates (input) and immunoprecipitates were analyzed by western blotting using specific antibodies. Densiometric quantification of CtIP-pS276 band in the GFP-Trap is shown (% indicate CtIP-pS276 band intensity vs CtIP band intensity). **(D)** Fork degradation was evaluated upon HU treatment in U2OS cells depleted of either endogenous CtIP or p38α. Box and whisker plots of IdU/CldU-tract length ratios for individual replication forks are shown. Numbers indicated above the individual plots represent the mean ratios ± standard deviation. Schematics of the CldU/IdU pulse-labelling protocol are shown (top). Western blotting of lysates from the same experiment is shown below. **(E)** Western blotting of lysates from cells used in figure 3E. **(F)** Western blotting of lysates from wild-type mouse embryonic fibroblasts (MEFs) and *Pin1^-/-^* MEFs.

**Figure S4.**
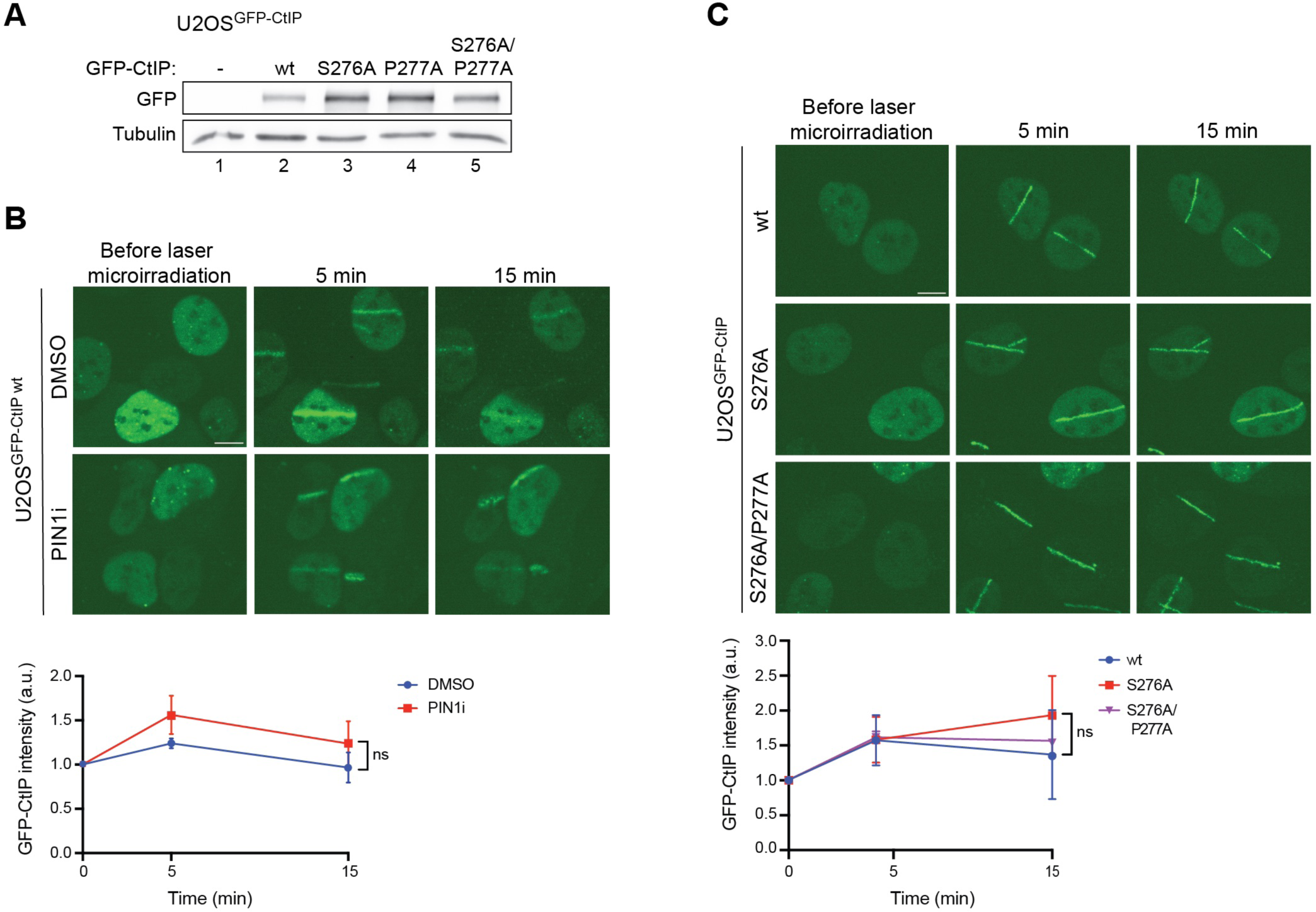
**(Related to Figure 4). (A)** Western blotting of lysates from U2OS cells inducibly expressing indicated GFP-CtIP variants and depleted of endogenous CtIP as employed in the SIRF analysis of figure 4C. **(B)** Laser micro-irradiation was performed in U2OS cells inducibly expressing GFP-CtIP wt treated with the PIN1 inhibitor (10 µM, 2h prior to laser micro-irradiation). Cells were grown in the presence of 5’-bromo-2’-deoxyuridine (BrdU) for 24h before micro-irradiation. Bottom: graph depicts GFP-CtIP intensity normalized on GFP pre-irradiation levels. Data are shown as mean ± standard deviation (n = 3). Representative images are shown (top, scale bars, 10 μm). **(C)** Laser micro-irradiation was performed in U2OS cells inducibly expressing GFP-CtIP wt, S276A and S276A/P277A *trans*-locked mutant depleted for endogenous CtIP. The next day cells were grown in the presence of BrdU for 24h before micro-irradiation. Two time points 5 and 15 minutes were taken after laser beam irradiation in live cell imaging. Bottom: Graph depicts GFP-CtIP intensity normalized on GFP pre-irradiation levels. Data are shown as mean ± standard deviation (n = 3). Representative images are shown (top, scale bars, 10 μm).

**Figure S5.**
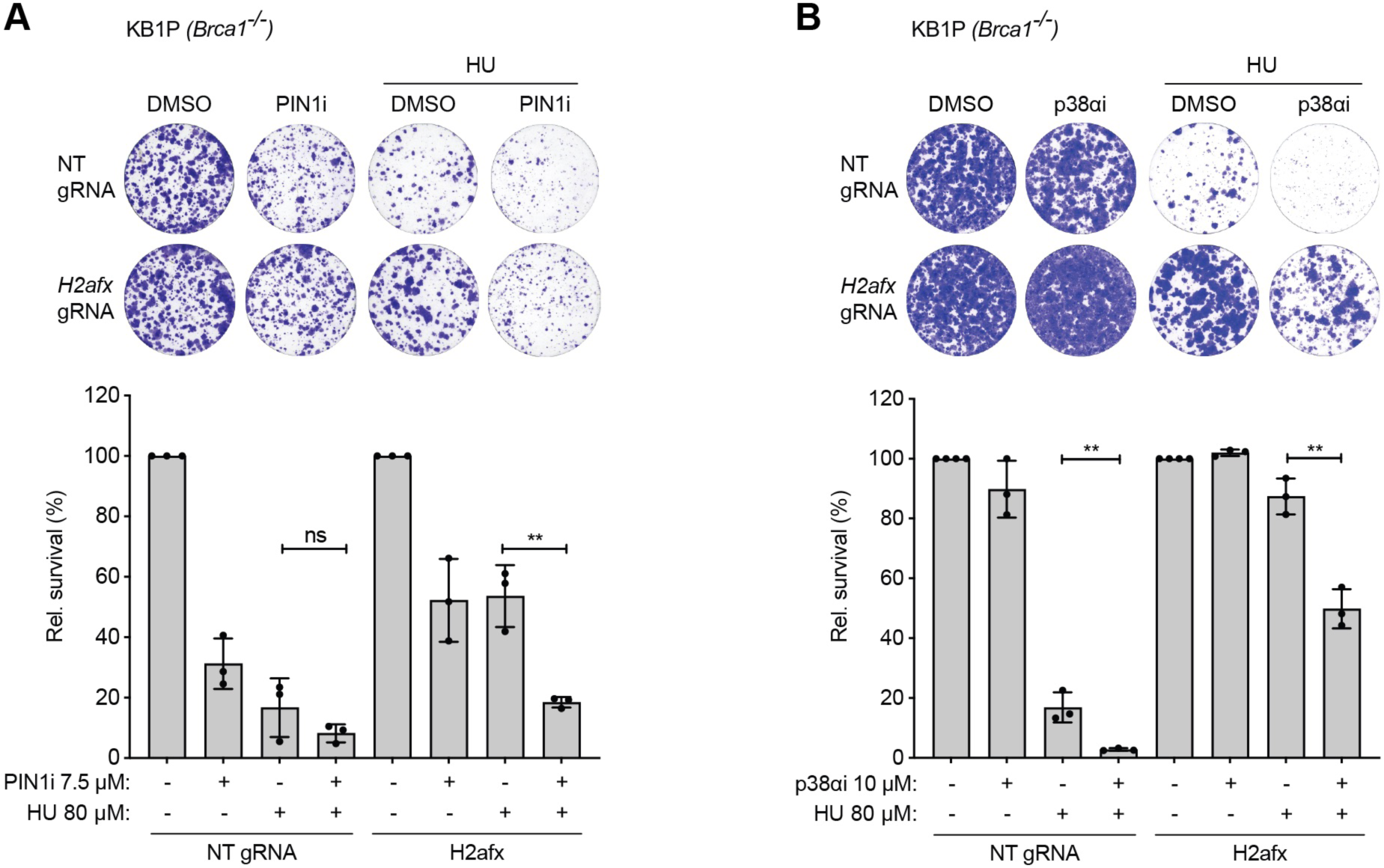
**(Related to Figure 5). (A)** Colony formation assay was performed in KB1P-derived *Trp53^-/-^*; *Brca1^-/-^* and *Trp53^-/-^*; *Brca1^-/-^*; *H2afx^-/-^* cells, either mock-treated or treated with the PIN1 inhibitor KPT-6566 (7.5 μM) and HU (80 μM) for 10 days. **(B)** Colony formation assay was performed in same cells as in (A), either mock-treated or treated with the p38α inhibitor PH-797804 (10 μM) and HU (80 μM) for 10 days. **(A and B)** Plotted values are mean ± standard deviation of three biological replicates. Representative images are shown (top).

## Bibliography

1. Zeman, M.K., and Cimprich, K.A. (2014). Causes and consequences of replication stress. Nat. Cell Biol. 16, 2–9. 10.1038/ncb2897.

2. Toledo, L.I., Altmeyer, M., Rask, M.-B., Lukas, C., Larsen, D.H., Povlsen, L.K., Bekker-Jensen, S., Mailand, N., Bartek, J., and Lukas, J. (2013). ATR Prohibits Replication Catastrophe by Preventing Global Exhaustion of RPA. Cell 155, 1088–1103. 10.1016/j.cell.2013.10.043.

3. Schlacher, K., Christ, N., Siaud, N., Egashira, A., Wu, H., and Jasin, M. (2011). Double-Strand Break Repair-Independent Role for BRCA2 in Blocking Stalled Replication Fork Degradation by MRE11. Cell 145, 529–542. 10.1016/j.cell.2011.03.041.

4. Przetocka, S., Porro, A., Bolck, H.A., Walker, C., Lezaja, A., Trenner, A., von Aesch, C., Himmels, S.-F., D’Andrea, A.D., Ceccaldi, R., et al. (2018). CtIP-Mediated Fork Protection Synergizes with BRCA1 to Suppress Genomic Instability upon DNA Replication Stress. Mol. Cell 72, 568–582.e6. 10.1016/j.molcel.2018.09.014.

5. Sartori, A.A., Lukas, C., Coates, J., Mistrik, M., Fu, S., Bartek, J., Baer, R., Lukas, J., and Jackson, S.P. (2007). Human CtIP promotes DNA end resection. Nature 450, 509–514. 10.1038/nature06337.

6. Huertas, P., and Jackson, S.P. (2009). Human CtIP Mediates Cell Cycle Control of DNA End Resection and Double Strand Break Repair. J. Biol. Chem. 284, 9558–9565. 10.1074/jbc.M808906200.

7. Huertas, P., Cortés-Ledesma, F., Sartori, A.A., Aguilera, A., and Jackson, S.P. (2008). CDK targets Sae2 to control DNA-end resection and homologous recombination. Nature 455, 689–692. 10.1038/nature07215.

8. Yu, X., and Chen, J. (2004). DNA Damage-Induced Cell Cycle Checkpoint Control Requires CtIP, a Phosphorylation-Dependent Binding Partner of BRCA1 C-Terminal Domains. Mol. Cell. Biol. 24, 9478–9486. 10.1128/MCB.24.21.9478-9486.2004.

9. Yu, J.H., Im, C.Y., and Min, S.-H. (2020). Function of PIN1 in Cancer Development and Its Inhibitors as Cancer Therapeutics. Front. Cell Dev. Biol. 8. 10.3389/fcell.2020.00120.

10. Fagiani, F., Govoni, S., Racchi, M., and Lanni, C. (2021). The Peptidyl-prolyl Isomerase Pin1 in Neuronal Signaling: from Neurodevelopment to Neurodegeneration. Mol. Neurobiol. 58, 1062–1073. 10.1007/s12035-020-02179-8.

11. Steger, M., Murina, O., Hühn, D., Ferretti, L.P., Walser, R., Hänggi, K., Lafranchi, L., Neugebauer, C., Paliwal, S., Janscak, P., et al. (2013). Prolyl Isomerase PIN1 Regulates DNA Double-Strand Break Repair by Counteracting DNA End Resection. Mol. Cell 50, 333–343. 10.1016/j.molcel.2013.03.023.

12. Reinhardt, H.C., Aslanian, A.S., Lees, J.A., and Yaffe, M.B. (2007). p53-Deficient Cells Rely on ATM- and ATR-Mediated Checkpoint Signaling through the p38MAPK/MK2 Pathway for Survival after DNA Damage. Cancer Cell 11, 175–189. 10.1016/j.ccr.2006.11.024.

13. Manke, I.A., Nguyen, A., Lim, D., Stewart, M.Q., Elia, A.E.H., and Yaffe, M.B. (2005). MAPKAP Kinase-2 Is a Cell Cycle Checkpoint Kinase that Regulates the G2/M Transition and S Phase Progression in Response to UV Irradiation. Mol. Cell 17, 37–48. 10.1016/j.molcel.2004.11.021.

14. Cánovas, B., Igea, A., Sartori, A.A., Gomis, R.R., Paull, T.T., Isoda, M., Pérez-Montoyo, H., Serra, V., González-Suárez, E., Stracker, T.H., et al. (2018). Targeting p38α Increases DNA Damage, Chromosome Instability, and the Anti-tumoral Response to Taxanes in Breast Cancer Cells. Cancer Cell 33, 1094–1110.e8. 10.1016/j.ccell.2018.04.010.

15. Campaner, E., Rustighi, A., Zannini, A., Cristiani, A., Piazza, S., Ciani, Y., Kalid, O., Golan, G., Baloglu, E., Shacham, S., et al. (2017). A covalent PIN1 inhibitor selectively targets cancer cells by a dual mechanism of action. Nat. Commun. 8, 15772. 10.1038/ncomms15772.

16. Daza-Martin, M., Starowicz, K., Jamshad, M., Tye, S., Ronson, G.E., MacKay, H.L., Chauhan, A.S., Walker, A.K., Stone, H.R., Beesley, J.F.J., et al. (2019). Isomerization of BRCA1–BARD1 promotes replication fork protection. Nature 571, 521–527. 10.1038/s41586-019-1363-4.

17. Dupré, A., Boyer-Chatenet, L., Sattler, R.M., Modi, A.P., Lee, J.-H., Nicolette, M.L., Kopelovich, L., Jasin, M., Baer, R., Paull, T.T., et al. (2008). A forward chemical genetic screen reveals an inhibitor of the Mre11–Rad50–Nbs1 complex. Nat. Chem. Biol. 4, 119–125. 10.1038/nchembio.63.

18. Kumar, S., Peng, X., Daley, J., Yang, L., Shen, J., Nguyen, N., Bae, G., Niu, H., Peng, Y., Hsieh, H.-J., et al. (2017). Inhibition of DNA2 nuclease as a therapeutic strategy targeting replication stress in cancer cells. Oncogenesis 6, e319–e319. 10.1038/oncsis.2017.15.

19. Luo, M.-L., Zheng, F., Chen, W., Liang, Z.-M., Chandramouly, G., Tan, J., Willis, N.A., Chen, C.-H., Taveira, M. de O., Zhou, X.Z., et al. (2020). Inactivation of the Prolyl Isomerase Pin1 Sensitizes BRCA1-Proficient Breast Cancer to PARP Inhibition. Cancer Res. 80, 3033–3045. 10.1158/0008-5472.CAN-19-2739.

20. Anand, R., Ranjha, L., Cannavo, E., and Cejka, P. (2016). Phosphorylated CtIP Functions as a Co-factor of the MRE11-RAD50-NBS1 Endonuclease in DNA End Resection. Mol. Cell 64, 940–950. 10.1016/j.molcel.2016.10.017.

21. Florensa, R., Bachs, O., and Agell, N. (2003). ATM/ATR-independent inhibition of cyclin B accumulation in response to hydroxyurea in nontransformed cell lines is altered in tumour cell lines. Oncogene 22, 8283–8292. 10.1038/sj.onc.1207159.

22. Rodríguez-Bravo, V., Guaita-Esteruelas, S., Salvador, N., Bachs, O., and Agell, N. (2007). Different S/M Checkpoint Responses of Tumor and Non–Tumor Cell Lines to DNA Replication Inhibition. Cancer Res. 67, 11648–11656. 10.1158/0008-5472.CAN-07-3100.

23. Llopis, A., Salvador, N., Ercilla, A., Guaita-Esteruelas, S., Barrantes, I. del B., Gupta, J., Gaestel, M., Davis, R.J., Nebreda, A.R., and Agell, N. (2012). The stress-activated protein kinases p38α/β and JNK1/2 cooperate with Chk1 to inhibit mitotic entry upon DNA replication arrest. Cell Cycle 11, 3627–3637. 10.4161/cc.21917.

24. Selness, S.R., Devraj, R.V., Devadas, B., Walker, J.K., Boehm, T.L., Durley, R.C., Shieh, H., Xing, L., Rucker, P.V., Jerome, K.D., et al. (2011). Discovery of PH-797804, a highly selective and potent inhibitor of p38 MAP kinase. Bioorg. Med. Chem. Lett. 21, 4066–4071. 10.1016/j.bmcl.2011.04.121.

25. Cubillos-Rojas, M., Loren, G., Hakim, Y.Z., Verdaguer, X., Riera, A., and Nebreda, A.R. (2023). Synthesis and Biological Activity of a VHL-Based PROTAC Specific for p38α. Cancers 15, 611. 10.3390/cancers15030611.

26. Taglialatela, A., Alvarez, S., Leuzzi, G., Sannino, V., Ranjha, L., Huang, J.-W., Madubata, C., Anand, R., Levy, B., Rabadan, R., et al. (2017). Restoration of replication fork stability in BRCA1- and BRCA2-deficient cells by inactivation of SNF2-family fork remodelers. Mol. Cell 68, 414–430.e8. 10.1016/j.molcel.2017.09.036.

27. Roy, S., Luzwick, J.W., and Schlacher, K. (2018). SIRF: Quantitative in situ analysis of protein interactions at DNA replication forks. J. Cell Biol. 217, 1521–1536. 10.1083/jcb.201709121.

28. Dungrawala, H., Rose, K.L., Bhat, K.P., Mohni, K.N., Glick, G.G., Couch, F.B., and Cortez, D. (2015). The Replication Checkpoint Prevents Two Types of Fork Collapse without Regulating Replisome Stability. Mol. Cell 59, 998–1010. 10.1016/j.molcel.2015.07.030.

29. Dibitetto, D., Liptay, M., Vivalda, F., Dogan, H., Gogola, E., González Fernández, M., Duarte, A., Schmid, J.A., Decollogny, M., Francica, P., et al. (2024). H2AX promotes replication fork degradation and chemosensitivity in BRCA-deficient tumours. Nat. Commun. 15, 4430. 10.1038/s41467-024-48715-1.

30. Celeste, A., Petersen, S., Romanienko, P.J., Fernandez-Capetillo, O., Chen, H.T., Sedelnikova, O.A., Reina-San-Martin, B., Coppola, V., Meffre, E., Difilippantonio, M.J., et al. (2002). Genomic Instability in Mice Lacking Histone H2AX. Science 296, 922–927. 10.1126/science.1069398.

31. Dias, M.P., Moser, S.C., Ganesan, S., and Jonkers, J. (2021). Understanding and overcoming resistance to PARP inhibitors in cancer therapy. Nat. Rev. Clin. Oncol. 18, 773–791. 10.1038/s41571-021-00532-x.

32. Canovas, B., and Nebreda, A.R. (2021). Diversity and versatility of p38 kinase signalling in health and disease. Nat. Rev. Mol. Cell Biol. 22, 346–366. 10.1038/s41580-020-00322-w.

33. Borisova, M.E., Voigt, A., Tollenaere, M.A.X., Sahu, S.K., Juretschke, T., Kreim, N., Mailand, N., Choudhary, C., Bekker-Jensen, S., Akutsu, M., et al. (2018). p38-MK2 signaling axis regulates RNA metabolism after UV-light-induced DNA damage. Nat. Commun. 9, 1017. 10.1038/s41467-018-03417-3.

34. Kuster, A., Mozaffari, N.L., Wilkinson, O.J., Wojtaszek, J.L., Zurfluh, C., Przetocka, S., Zyla, D., von Aesch, C., Dillingham, M.S., Williams, R.S., et al. (2021). A stapled peptide mimetic of the CtIP tetramerization motif interferes with double-strand break repair and replication fork protection. Sci. Adv. 7, eabc6381. 10.1126/sciadv.abc6381.

35. Ray Chaudhuri, A., Callen, E., Ding, X., Gogola, E., Duarte, A.A., Lee, J.-E., Wong, N., Lafarga, V., Calvo, J.A., Panzarino, N.J., et al. (2016). Replication fork stability confers chemoresistance in BRCA-deficient cells. Nature 535, 382–387. 10.1038/nature18325.

36. Koikawa, K., Kibe, S., Suizu, F., Sekino, N., Kim, N., Manz, T.D., Pinch, B.J., Akshinthala, D., Verma, A., Gaglia, G., et al. (2021). Targeting Pin1 renders pancreatic cancer eradicable by synergizing with immunochemotherapy. Cell 184, 4753–4771.e27. 10.1016/j.cell.2021.07.020.

37. Chen, L., Xu, X., Wen, X., Xu, S., Wang, L., Lu, W., Jiang, M., Huang, J., Yang, D., Wang, J., et al. (2019). Targeting PIN1 exerts potent antitumor activity in pancreatic ductal carcinoma via inhibiting tumor metastasis. Cancer Sci. 110, 2442–2455. 10.1111/cas.14085.

38. Yang, L., Sun, X., Ye, Y., Lu, Y., Zuo, J., Liu, W., Elcock, A., and Zhu, S. (2019). p38α Mitogen-Activated Protein Kinase Is a Druggable Target in Pancreatic Adenocarcinoma. Front. Oncol. 9. 10.3389/fonc.2019.01294.

39. Alam, M.S., Gaida, M.M., Bergmann, F., Lasitschka, F., Giese, T., Giese, N.A., Hackert, T., Hinz, U., Hussain, S.P., Kozlov, S.V., et al. (2015). Selective inhibition of the p38 alternative activation pathway in infiltrating T cells inhibits pancreatic cancer progression. Nat. Med. 21, 1337–1343. 10.1038/nm.3957.

40. Singh, S.P., Dosch, A.R., Mehra, S., De Castro Silva, I., Bianchi, A., Garrido, V.T., Zhou, Z., Adams, A., Amirian, H., Box, E.W., et al. (2024). Tumor Cell–Intrinsic p38 MAPK Signaling Promotes IL1α-Mediated Stromal Inflammation and Therapeutic Resistance in Pancreatic Cancer. Cancer Res. 84, 1320–1332. 10.1158/0008-5472.CAN-23-1200.

41. Hoadley, K.A., Yau, C., Hinoue, T., Wolf, D.M., Lazar, A.J., Drill, E., Shen, R., Taylor, A.M., Cherniack, A.D., Thorsson, V., et al. (2018). Cell-of-Origin Patterns Dominate the Molecular Classification of 10,000 Tumors from 33 Types of Cancer. Cell 173, 291–304.e6. 10.1016/j.cell.2018.03.022.

42. Witkiewicz, A.K., McMillan, E.A., Balaji, U., Baek, G., Lin, W.-C., Mansour, J., Mollaee, M., Wagner, K.-U., Koduru, P., Yopp, A., et al. (2015). Whole-exome sequencing of pancreatic cancer defines genetic diversity and therapeutic targets. Nat. Commun. 6, 6744. 10.1038/ncomms7744.

43. Waters, A.M., and Der, C.J. (2018). KRAS: The Critical Driver and Therapeutic Target for Pancreatic Cancer. Cold Spring Harb. Perspect. Med. 8, a031435. 10.1101/cshperspect.a031435.

44. Ferretti, L.P., Himmels, S.-F., Trenner, A., Walker, C., Aesch, C. von, Eggenschwiler, A., Murina, O., Enchev, R.I., Peter, M., Freire, R., et al. (2016). Cullin3-KLHL15 ubiquitin ligase mediates CtIP protein turnover to fine-tune DNA-end resection. Nat. Commun. 7, 1–16. 10.1038/ncomms12628.

45. Murina, O., von Aesch, C., Karakus, U., Ferretti, L.P., Bolck, H.A., Hänggi, K., and Sartori, A.A. (2014). FANCD2 and CtIP Cooperate to Repair DNA Interstrand Crosslinks. Cell Rep. 7, 1030–1038. 10.1016/j.celrep.2014.03.069.

46. Davies, O.R., Forment, J.V., Sun, M., Belotserkovskaya, R., Coates, J., Galanty, Y., Demir, M., Morton, C.R., Rzechorzek, N.J., Jackson, S.P., et al. (2015). CtIP tetramer assembly is required for DNA-end resection and repair. Nat. Struct. Mol. Biol. 22, 150–157. 10.1038/nsmb.2937.

47. Merrick, C.J., Jackson, D., and Diffley, J.F.X. (2004). Visualization of Altered Replication Dynamics after DNA Damage in Human Cells*. J. Biol. Chem. 279, 20067–20075. 10.1074/jbc.M400022200.

48. Nieminuszczy, J., Schwab, R.A., and Niedzwiedz, W. (2016). The DNA fibre technique – tracking helicases at work. Methods 108, 92–98. 10.1016/j.ymeth.2016.04.019.

49. Schindelin, J., Arganda-Carreras, I., Frise, E., Kaynig, V., Longair, M., Pietzsch, T., Preibisch, S., Rueden, C., Saalfeld, S., Schmid, B., et al. (2012). Fiji: an open-source platform for biological-image analysis. Nat. Methods 9, 676–682. 10.1038/nmeth.2019.

50. Porro, A., Berti, M., Pizzolato, J., Bologna, S., Kaden, S., Saxer, A., Ma, Y., Nagasawa, K., Sartori, A.A., and Jiricny, J. (2017). FAN1 interaction with ubiquitylated PCNA alleviates replication stress and preserves genomic integrity independently of BRCA2. Nat. Commun. 8, 1073. 10.1038/s41467-017-01074-6.

51. Krajewska, M., Fehrmann, R.S.N., de Vries, E.G.E., and van Vugt, M.A.T.M. (2015). Regulators of homologous recombination repair as novel targets for cancer treatment. Front. Genet. 6. 10.3389/fgene.2015.00096.

52. Ceppi, I., Cannavo, E., Bret, H., Camarillo, R., Vivalda, F., Thakur, R.S., Romero-Franco, A., Sartori, A.A., Huertas, P., Guérois, R., et al. (2023). PLK1 regulates CtIP and DNA2 interplay in long-range DNA end resection. Genes Dev. 37, 119–135. 10.1101/gad.349981.122.

53. Cannavo, E., and Cejka, P. (2014). Sae2 promotes dsDNA endonuclease activity within Mre11–Rad50–Xrs2 to resect DNA breaks. Nature 514, 122–125. 10.1038/nature13771.

54. Teloni, F., Michelena, J., Lezaja, A., Kilic, S., Ambrosi, C., Menon, S., Dobrovolna, J., Imhof, R., Janscak, P., Baubec, T., et al. (2019). Efficient Pre-mRNA Cleavage Prevents Replication-Stress-Associated Genome Instability. Mol. Cell 73, 670–683.e12. 10.1016/j.molcel.2018.11.036.

55. Johnson, J.L., Yaron, T.M., Huntsman, E.M., Kerelsky, A., Song, J., Regev, A., Lin, T.-Y., Liberatore, K., Cizin, D.M., Cohen, B.M., et al. (2023). An atlas of substrate specificities for the human serine/threonine kinome. Nature 613, 759–766. 10.1038/s41586-022-05575-3.

